# The RNA splicing factor PRPF8 is required for left-right organiser cilia differentiation and determination of cardiac left-right asymmetry via regulation of *Arl13b* splicing

**DOI:** 10.1101/2025.05.22.654869

**Authors:** Fangfei Jiang, Michael Boylan, Dale W. Maxwell, Wasay Mohiuddin Shaikh Qureshi, Charlie F. Rowlands, Gennadiy Tenin, Karen Mitchell, Louise A. Stephen, Elton J. R. Vasconcelos, Dapeng Wang, Tong Chen, Junzhe Zha, Jingshu Liu, Nouf Althali, Dragos V. Leordean, Meurig T. Gallagher, Basudha Basu, Katarzyna Szymanska, Advait Veeraghanta, Bernard Keavney, Martin J. Humphries, Jamie Ellingford, David Smith, Colin A. Johnson, Raymond T. O’Keefe, Sudipto Roy, Kathryn E. Hentges

## Abstract

Cilia function in the left-right organizer (LRO) is critical for determining internal organ asymmetry in vertebrates. To further understand the genetics of left-right asymmetry, we isolated a mouse mutant with laterality defects, *l11Jus27,* from a random mutagenesis screen. *l11Jus27* mutants carry a missense mutation in the pre-mRNA processing factor, *Prpf8*. *cephalophŏnus (cph)* mutant zebrafish, carrying a protein truncating mutation in *prpf8*, phenocopy the laterality defects of *l11Jus27* mutants. *Prpf8* mutant mouse and fish embryos have increased expression of an alternative transcript encoding the cilium-associated protein, ARL13B, that lacks exon 9. In zebrafish, over-expression of the *arl13b* transcript lacking exon 9 perturbed cilium formation and caused laterality defects. The shorter ARL13B protein isoform lacked interactions with intraflagellar transport proteins. Our data suggest that PRPF8 plays a prominent role in LRO cilia by through the regulation of alternative splicing of ARL13B, thus uncovering a new mechanism for cilia-linked developmental defects.

## Introduction

Cilia are specialised hair-like organelles required for many cellular functions (1). Motile cilia produce a rhythmic waving or beating motion, whereas non-motile cilia act as sensory antennae for cells (1, 2). Diseases caused by cilia dysfunction are collectively known as ciliopathies. Mutations in the wide range of genes required for cilia function cause ciliopathies, with most phenotypes displaying both genetic heterogeneity and variable expressivity (3). In humans, mutations in several spliceosomal pre-mRNA processing factor proteins cause autosomal dominant Retinitis Pigmentosa (RP) (4–6), a ciliopathy causing progressive retinal atrophy and loss of sight with age. RP-causing mutations in Pre-mRNA Processing Factor 8 (PRPF8) produce multiple splicing defects (5, 7–11). However, PRPF8 may have a more direct role in cilia functionality. PRPF8 localises to the basal body of primary cilia, and knocking down PRPF8 in human and mouse cells (12, 13), zebrafish embryos (13) and *C. elegans* (14), inhibits ciliogenesis.

In vertebrate embryogenesis, specialised motile cilia in the LRO play a critical role in left-right axis specification (2, 15). In mice, laterality is established at the ventral node (2), the mammalian LRO. The node is a teardrop shaped pit at the posterior end of the notochord with motile cilia; these cilia generate a leftward flow of fluid within the node cavity thought to be detected by immotile cilia on adjacent perinodal ‘crown’ cells (16–20). In response, expression of the genes *Cerl2* and *Nodal*, becomes restricted preferentially on the right and left side of the node, respectively (21–23). Nodal in the left lateral plate mesoderm (LPM) induces expression of the Nodal inhibitors *Lefty1* and *Lefty2* (24, 25) at the midline and left LPM, respectively (26). Nodal in the LPM also induces expression of the transcription factor *Pitx2* (27). *Pitx2* is considered to be the effector of Nodal signalling, and is required for the correct positioning and morphogenesis of organs including the heart, lungs, gut and blood vessels (28–30), but is dispensable for heart looping directionality (reviewed in (31)).

A recessive lethal mouse mutant with laterality defects was isolated from a random chemical mutagenesis screen: the *l11Jus27* mouse (32, 33). The causative mutation in this mouse is an asparagine (N) to serine (S) missense mutation at residue 1531 in *Prpf8*, the mouse orthologue of human *PRPF8*. *Prpf8^N1531S/N1531S^* embryos (hereafter referred to as *Prpf8^N1531S^*mutants or *Prpf8^N1531S^* hom) have impaired LRO cilia formation and motility. Zebrafish *cph* mutants (carrying a Prpf8 truncation mutation) phenocopy laterality defects seen in *Prpf8^N1531S^* mutant mouse embryos. Moreover, modelling the analogous *Prpf8^N1531S^* mutation in yeast resulted in splicing defects. Bulk RNA-seq analysis of *Prpf8^N1531S^* mutant mouse embryos showed increased skipping of exon 9 in the transcript of the Arf-like GTPase *Arl13b;* the ARL13B protein is specifically localised to the ciliary membrane (34, 35) where it functions in regulating cilia length, and thus impacts upon beat pattern and flow (36–38), as well as ciliary trafficking via interactions with intraflagellar transport (IFT) proteins (39, 40). Importantly, over-expression of the shorter a*rl13b* transcript (lacking exon 9) in wild type zebrafish re-capitulated many aspects of the *Prpf8^N1531S^*mutant phenotype. Additionally, the ARL13B protein lacking the region encoded by exon 9 showed reduced binding to IFT partners. Combined, these results suggest that altering *Arl13b* alternative transcript ratios may be the mechanistic basis for the laterality defects observed in *Prpf8* mutants. The data presented here reveal specific functions for PRPF8 in cilia function and embryonic patterning, which may also provide new insights into the role of PRPF8 in disease pathology in patients with RP.

## Results

### Phenotypic analysis of *l11Jus27* mouse mutants

The *l11Jus27* mouse line was isolated from a random chemical mutagenesis screen, which employed a balancer chromosome to facilitate mapping of mutations (32). *l11Jus27* heterozygotes are viable and fertile. At E8.25, *l11Jus27* homozygous embryos appear grossly normal (Supplemental Fig. 1a-b) but are developmentally delayed (Supplemental Fig. 1c). At E9.5 the *l11Jus27* homozygotes have an open neural tube, distended heart tube, and a high incidence of reversed cardiac looping (Fig. 1a). At E10.5 less severely affected mutants have milder cardiac defects and developmental delay (Supplemental Fig. 1d-f). More severely delayed embryos have also failed to undergo chorioallantoic fusion (Supplemental Fig. 1f). *l11Jus27* homozygous embryos have impaired yolk sac vascularisation at E9.5 (Supplemental Fig. 1g-h) and frequently do not have patent umbilical vessels (Supplemental Fig 1i-j), probably leading to the failure in chorioallantoic fusion observed in mutant embryos at E10.5 (Supplemental Fig 1j; penetrance reported in Supplemental Table 1). The majority of homozygous mutant embryos died by E11.5, and none were isolated after E12.5.

**Fig 1.**
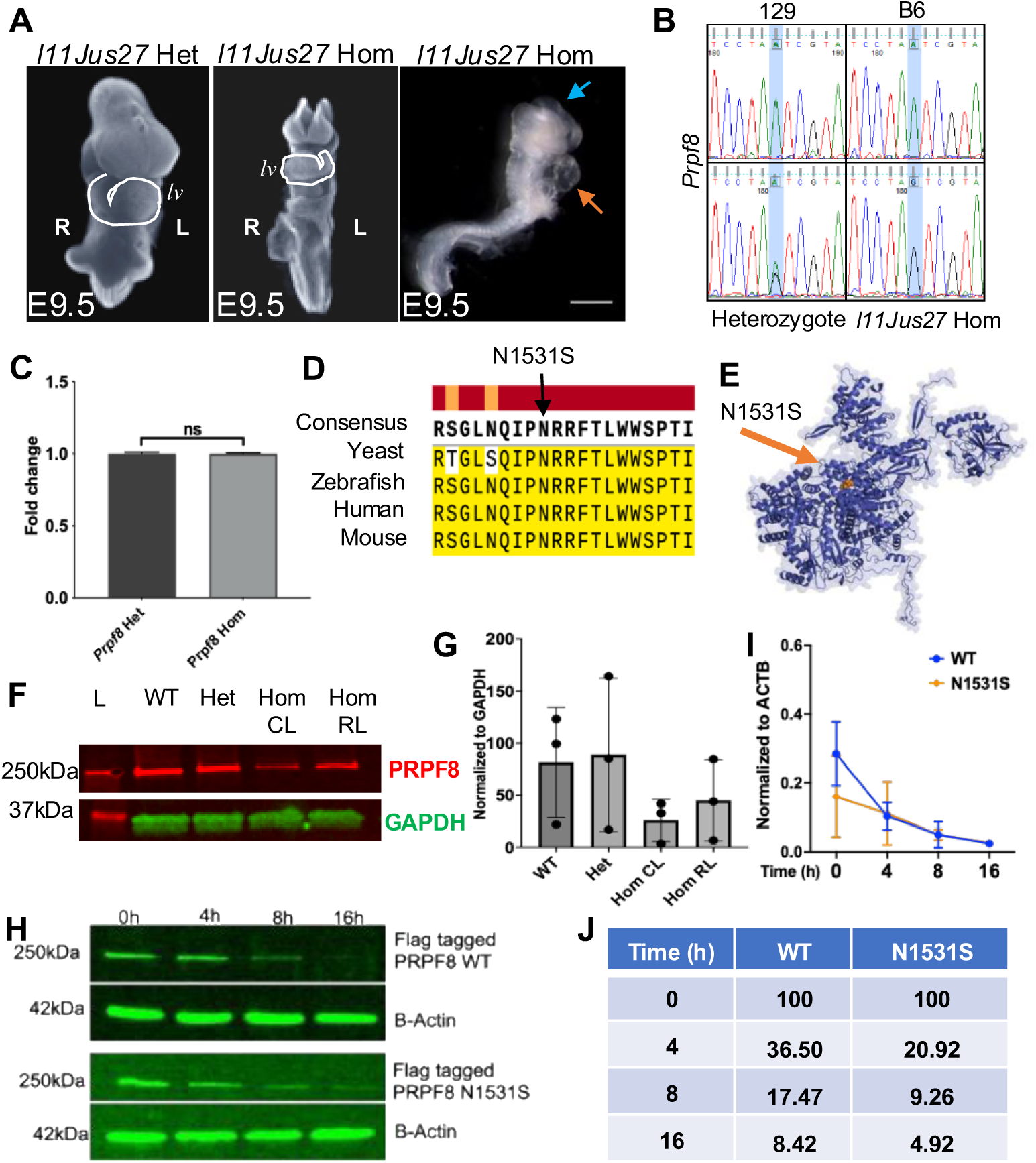
Genetic analysis of *l11Jus27* mice. A) Propidium iodide stained E9.5 *l11Jus27* heterozygote showing correct leftward looping of heart (heart tube outlined in white; lv= left ventricle), *l11Jus27* homozygote showing reversed looping of heart (heart tube outlined in white; lv = morphological left ventricle), *l11Jus27* homozygote showing open neural tube (blue arrow) and distended heart tube (orange arrow). B) Sanger sequencing of individual mouse embryos confirming that the N1531S substitution is present in heterozygous embryos and homozygous embryos carrying the *l11Jus27* phenotype, but not in wild type mice from either a 129S5 (129) or C57BL/6 (B6) background. C) qPCR quantification of *Prpf8* expression levels in *Prpf8^N1531S^* heterozygous and homozygous mutant whole embryos at E10.5. Results from the average of three biological and three technical replicates are presented. Expression levels were normalised to *Gapdh* expression. D) Clustal analysis of PRPF8 protein orthologues in mouse, zebrafish, human and *Saccharomyces cerevisiae* showing that residue 1531 is conserved across multiple eukaryotes. Identical residues are highlighted in yellow. E) Protein structure of PRPF8 with amino acid replacement at residue 1531 shown in red. F) Western blot confirms PRPF8 protein is present in *Prpf8^N1531S^* heterozygotes and homozygous mutant whole embryos with either correct heart looping (CL) or reversed heart looping (RL) at E10.5. G) Quantification of PRPF8 protein levels from Western blots. Each sample was normalised to GAPDH protein expression levels. H) Western blot analysis of mammalian cells transfected with epitope tagged PRPF8 constructs at specified time points after cycloheximide treatment. I) Quantification of PRPF8 expression levels normalised to ACTB for wild type and PRPF8^N1531S^ alleles following cycloheximide treatment. J) PRPF8 WT and N1531S protein half-life percentage decrease as compared to protein levels present at 0h. Numbers report percentage of original protein remaining at each time point. Differences in degradation rate are non-significant (Wilcoxon test p=0.125).

### The *l11Jus27* line contains a *Prpf8^N1531S^* mutation

The *l11Jus27* mutant mouse line was isolated from a balancer chromosome mutagenesis screen; therefore the causative mutation was known to reside in a 24cM region of mouse chromosome 11 (32). Meiotic mapping further refined the *l11Jus27* candidate interval (Supplemental Fig. 1k) (41). Genome sequencing of an E10.5 homozygous mutant embryo revealed a novel point mutation of an A to G transition within the genetic candidate interval at position chr11: 75,391,978 (mm39), which produces a N1531S missense mutation in *Prpf8*. This sequence alteration was confirmed in an additional 3 homozygous mutant embryos and was not present in wild type C57BL/6 or 129S5 animals (Fig. 1b). Thus, we concluded the *l11Jus27* line carries a PRPF8 N1531S missense mutation, and *l11Jus27* homozygous embryos will now be referred to as *Prpf8* hom. We confirmed by quantitative RT-PCR that the N1531S mutation does not affect *Prpf8* transcript levels (Fig. 1c). PRPF8 is highly conserved (42), and residue N1531 is conserved between multiple eukaryotes (Fig. 1d). The N1531S amino acid substitution is in a linker region near the endonuclease-like domain (Fig. 1e; (43)) and is predicted to be damaging by PolyPhen and SIFT (44, 45) in both mouse and human proteins. At E10.5, mutants appeared to display a reduction in PRPF8 protein levels, but this was not significantly different (Fig. 1f-g; one-way ANOVA p=0.4212)(12). To test if the PRPF8 mutation altered protein stability, we expressed an N-terminal FLAG epitope-tagged PRPF8 construct in mammalian cells (Fig. 1h) and performed a cycloheximide assay to determine the PRPF8 degradation rate. We detected a reduced level of epitope tagged PRPF8^N1531S^ protein compared to wild type protein at each time point in the assay, but the overall degradation rate of the N1531S mutant was similar to the wild type construct (Wilcoxon test p=0.125; Fig. 1i-j), and thus did not indicate significant evidence of increased protein degradation.

### Nodal cilia are present but dysmotile in *Prpf8 ^N1531S^* mutants

Heart looping defects are associated with ciliopathies (46, 47). Scanning electron microscopy (SEM) revealed that control and *Prpf8^N1531S^* mutant embryos possessed abundant nodal cilia at the 2-3 somite stage (Fig. 2a-d). Control embryos had stereotypical pit shaped nodes (Fig. 2a, n=7), whereas *Prpf8^N1531S^* mutant embryonic nodes appeared unusually flat (Fig. 2b, *n*=6/8), suggesting that node morphogenesis is disrupted in *Prpf8^N1531S^* mutant embryos. While there was a trend towards fewer cilia in *Prpf8^N1531S^* mutant embryos, this trend was not statistically significant (Fig. 2e; unpaired t-test of late head fold (LHF) stage data, p=0.1934; for 2-3 somite data p=0.1494). Transmission electron microscopy (TEM) analysis of nodal cilia did not reveal any overt differences between homozygous *Prpf8^N1531S^* mutant embryos and controls (Supplemental Fig 2). Because mutations in the spliceosome have been found to affect the rate of cell division (48), we performed immunofluorescence staining for phospho-histone H3 to identify cells in mitosis. There was no difference in the percentage of dividing cells between genotypes (Fig. 2f; unpaired t-test p=0.7918). Using acetylated tubulin as a marker of cilia (Fig. 2g-h), we found that node cilia length in homozygous *Prpf8^N1531S^* mutant embryos was significantly shorter than in heterozygous control embryos (Fig. 2i, unpaired t-test p=0.0408).

**Fig 2.**
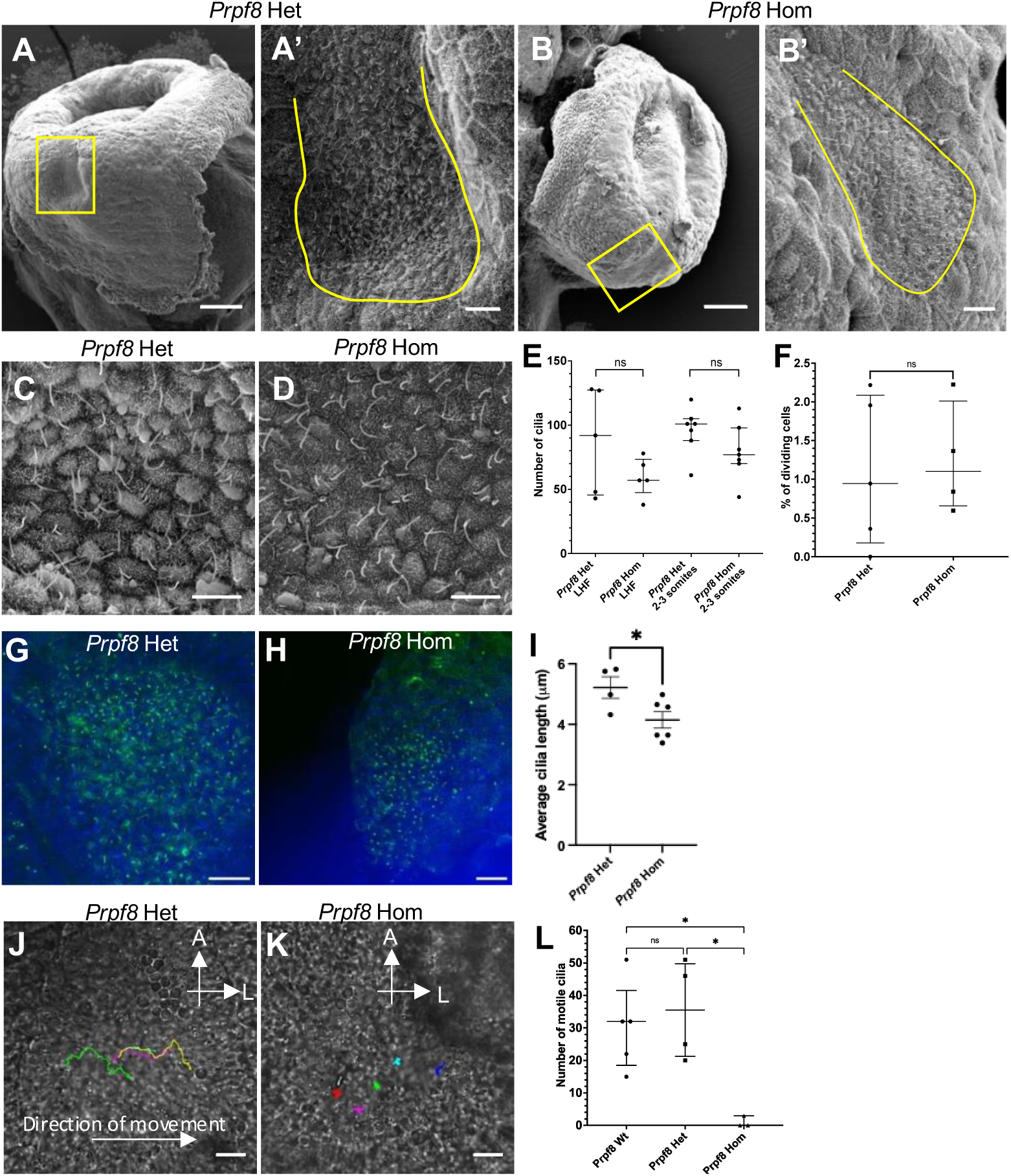
Analysis of *Prpf8^N1531S^* heterozygous and homozygous node cilia. A-D) SEM images of 2-3 somite stage *Prpf8^N1531S^* heterozygous (A, C) and homozygous (B, D) mouse embryos. A) The pit shape of the node (yellow boxes) is obvious in *Prpf8^N1531S^* heterozygotes. *Prpf8^N1531S^* mutant embryos (B) have a flat node. Scale bar = 100μm. A’, B’) Higher magnification images of boxed region in A and B. The node (yellow outline) is ciliated in both genotypes. Scale bar = 10um. C-D) Higher magnification images of the node at 2-3 somite stage. No gross morphological difference in cilia was observed between genotypes. Scale bar = 5μm. E) Heterozygotes and homozygotes have equivalent numbers of nodal cilia at the LHF and 2-3 somite stage (unpaired t-test). F) Cell proliferation was unaltered between genotypes (unpaired t-test). G-H) Mouse node cilia labelled with acetylated tubulin in heterozygous control (G) and *Prpf8^N1531S^* mutant embryos (H). Nuclei are stained with DAPI. I) Measurement of node cilia length reveals that *Prpf8^N1531S^* mutant cilia are significantly shorter than heterozygous controls (unpaired t-test p=0.0408). J-K) Tracking the travel over time of five microbeads in the node revealed directional movement in *Prpf8^N1531S^* heterozygotes (J) but only Brownian motion in *Prpf8^N1531S^* homozygotes (K). Direction of movement is indicated with an arrow. A= anterior, L = left. L) Quantification of motile nodal cilia in *Prpf8^+/+^*, *Prpf8^N1531S^* heterozygous and *Prpf8^N1531S^* homozygous embryos at the 2-3 somite stage revealed significant differences between the three genotypes (p=0.0058, Welch’s ANOVA). Post-hoc testing with Dunnett’s T3 test showed that the number of motile cilia in *Prpf8^N1531S^* homozygotes was significantly lower compared to *Prpf8^+/+^* (p=0.0222) and *Prpf8^N1531S/+^* embryos (p=0.0487), but not between *Prpf8^N1531S/+^*and *Prpf8^+/+^* embryos (p=0.9332).

LRO fluid flow generation is related to cilia length (38). Therefore, we evaluated whether flow was established correctly in *Prpf8^N1531S^* mutant embryo nodes. We placed microbeads in the node of embryos mounted in a chamber slide to detect the direction of nodal flow. While almost all heterozygotes established flow towards the left of the node (*n*=5/6; Fig. 2j; Supplemental Movie File 1), almost all *Prpf8^N1531S^* mutants failed to establish any directional fluid flow (*n*=5/6; Fig. 2k); instead the microbeads showed random Brownian motion (Supplemental Movie file 2). We next investigated whether cilia motility was impaired in mutant embryos. Directly visualising cilia movement using high-speed videomicroscopy revealed that wild type embryos had strongly beating cilia (*n*=5 Supplemental Movie File 3), as did *Prpf8^N1531S/+^* embryos (*n*=4/4). However, *Prpf8^N1531S^* homozygotes had very few motile cilia (*n*=3; Supplemental Movie File 4). We quantified the number of motile cilia (Fig. 2l) and found significant differences between the three genotypes (p=0.0058, Welch’s ANOVA). Posthoc testing revealed that the number of motile cilia in *Prpf8^N1531S^* homozygotes was significantly lower compared to *Prpf8^+/+^* (p=0.0222) and *Prpf8^N1531S/+^* embryos (p=0.0487), but not between *Prpf8^N1531S/+^* and *Prpf8^+/+^* embryos.

PRPF8 has been reported to localise to the base of cilia (12–14). To determine if the *Prpf8^N1531S^* missense mutation affected protein localisation, we used immunofluorescence to analyse node cilia. We confirmed that PRPF8 protein can be detected at the base of some cilia in heterozygous (Supplemental Fig. 3 a-c) and *Prpf8^N1531S^* homozygous mutant embryos within the node (Supplemental Fig. 3 g-i). A transverse projection of the Z-stack of confocal images confirmed that the node in heterozygous embryos formed a pit like structure (Supplemental 3d-f) but in homozygotes the node was flatter (Supplemental 3 j-l). These findings suggest that *Prpf8^N1531S^* homozygous mutants have defects in node structure, cilia motility, and nodal fluid flow that are consistent with downstream defects in L-R axis formation.

### Laterality gene expression analysis reveals pathway defects in ***Prpf8^N1531S^*** mutants

We next determined whether the L-R axis pathway was disturbed by the defects in the node. We performed whole-mount *in situ* hybridisation for several genes that are important for L-R axis formation and found that *Prpf8^N1531S^* mutant embryos had highly aberrant laterality gene expression patterns. *Shh*, needed for L-R axis establishment (49), was expressed in the midline and node of both control and *Prpf8^N1531S^* mutant embryos (Fig. 3a-b), though half of *Prpf8^N1531S^* mutants displayed fainter and discontinuous midline *Shh* expression (*n*=3/6) *Cerl2* (Fig. 3c-e) and *Nodal* (Fig. 3f-h) expression was highly variable, with normal, reversed, symmetric and absent expression patterns observed for both genes in the mutants. The LPM of *Prpf8^N1531S^*mutants usually lacked *Nodal* expression (*n*=19/21), and when *Nodal* was expressed, the expression domain did not include the anterior LPM. Accordingly, both *Lefty1* and *Lefty2* were not expressed in *Prpf8^N153S1^* mutant embryos (*n*=10/10) (Fig 3i-j). *Pitx2* expression in the LPM and subsequent secondary heart field (50), which is induced by Nodal activity, was variable, with normal (n=13/38), reversed (n=8/38), bilateral (n=8/38) and absent (n=9/38) expression in *Prpf8^N153S1^* mutant embryos (Fig 3k-p). *Pitx2* is normally expressed on the same side as the developing ventricle. We quantified both *Pitx2* expression sidedness and heart looping direction in mutant embryos (Fig 3q). In embryos where the sidedness of *Pitx2* expression was unambiguous, 29% (n=6/21) expressed *Pitx2* on the opposite side of the embryo from the inflow tract of the heart (Fig 3r). These data led us to hypothesise that the *Prpf8^N1531S^* mutants with reversed cardiac looping exhibited heterotaxy rather than *situs inversus.* To test this hypothesis, we examined the expression of *Barx1* in the developing stomach (51), which is found left of the embryonic midline. Both control and mutant embryos, including those with reversed cardiac looping (n=6), maintained *Barx1* expression left of the midline, (Fig 3s-t), supporting the hypothesis that *Prpf8^N1531S^* mutant embryos model heterotaxy, not *situs inversus*. The early lethality of *Prpf8^N1531S^* mutant embryos precluded examining the laterality of other organ systems.

**Fig 3.**
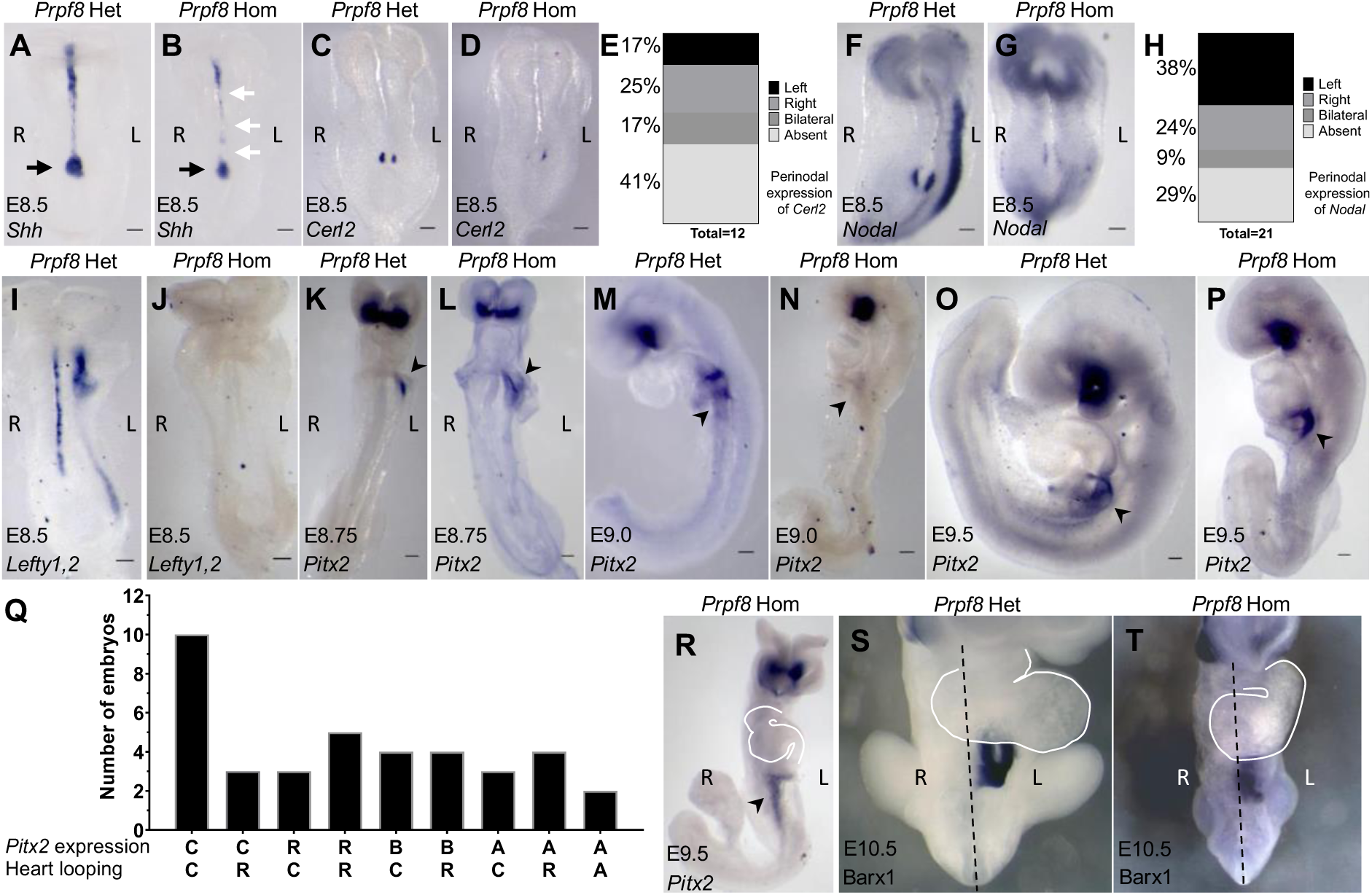
Wholemount *in situ* hybridisation expression analysis of laterality genes in *Prpf8^N1531S^* heterozygous and homozygous mutant embryos. A) *Prpf8^N1531S^*heterozygote showing *Shh* midline and node expression (black arrow). B) *Prpf8^N1531S^* homozygote showing intact *Shh* expression at the node (black arrow), but discontinuous midline staining (white arrows). C) E8.5 *Prpf8^N1531S^* heterozygote with upregulation of *Cerl2* on the right vs the left of the node. D) *Prpf8^N1531S^*homozygote with reversed *Cerl2* expression. E) Graphical representation of *Cerl2* expression patterns seen in *Prpf8^N1531S^*homozygotes revealing a high degree of variability. F) E8.5 *Prpf8^N1531S^* heterozygote showing *Nodal* expression pattern with upregulation left of the node and left LPM. G) *Prpf8^N1531S^* homozygote showing reduced and reversed perinodal *Nodal* expression and no expression in the LPM. H) Graphical representation of *Nodal* perinodal expression in *Prpf8^N1531S^* homozygotes revealing variable expression patterns. I) E8.5 *Prpf8^N1531S^* heterozygote showing *Lefty1, 2* expression along the midline and left LPM, detected with a probe complimentary to both mRNAs; no expression was detected in *Prpf8^N1531S^* homozygotes (J). K-P) *Pitx2* expression in the LPM (black arrowheads) at E8.75 (K,L), E9.0 (M,N) and E9.5 (O,P) with *Prpf8^N1531S^*heterozygotes showing left-sided *Pitx2* expression (K,M,O) and *Prpf8^N1531S^*homozygotes showing randomised Pitx2 expression (L,N,P). Q) Graphical representation of data comparing sidedness of LPM *Pitx2* expression with heart looping direction at E9.5 revealing a disconnect between expression sidedness (*Pitx2*) and heart looping (heart). C= correct, R= reversed, B= bilateral, A= absent. R) E9.5 expression of *Pitx2* in *Prpf8^N1531S^*homozygote showing discordance between heart looping and *Pitx2* expression sidedness (arrow). S,T) E10.5 expression of *Barx1* labelling the developing stomach in *Prpf8^N1531S^* heterozygote (S) and in *Prpf8^N1531S^* homozygote (T). Dashed line indicates the midline, and the heart is outlined in white. L= left, R= right. Scale bars = 0.5mm.

### Phenocopy of *Prpf8 ^N1531S^* laterality defects in *prpf8* mutant zebrafish

To evaluate if laterality defects present in the *Prpf8^N1531S^*mouse showed evolutionary conservation, we examined the ENU mutant zebrafish strain, *cph* (52). This mutant line carries an E396* substitution in Prpf8, predicting a severely truncated protein (52). Whereas wild type (Fig. 4a) and heterozygous embryos are phenotypically normal, *cph* homozygous mutant embryos displayed a curled down body axis and delayed heart looping (Fig. 4b). *cph* homozygotes displayed randomised cardiac looping: 27% of *cph* mutants had correctly looped, 42% unlooped, and 31% reversed looped hearts, compared to 97%, 1% and 2%, respectively, for wild type fish (Fig. 4c).

**Fig 4.**
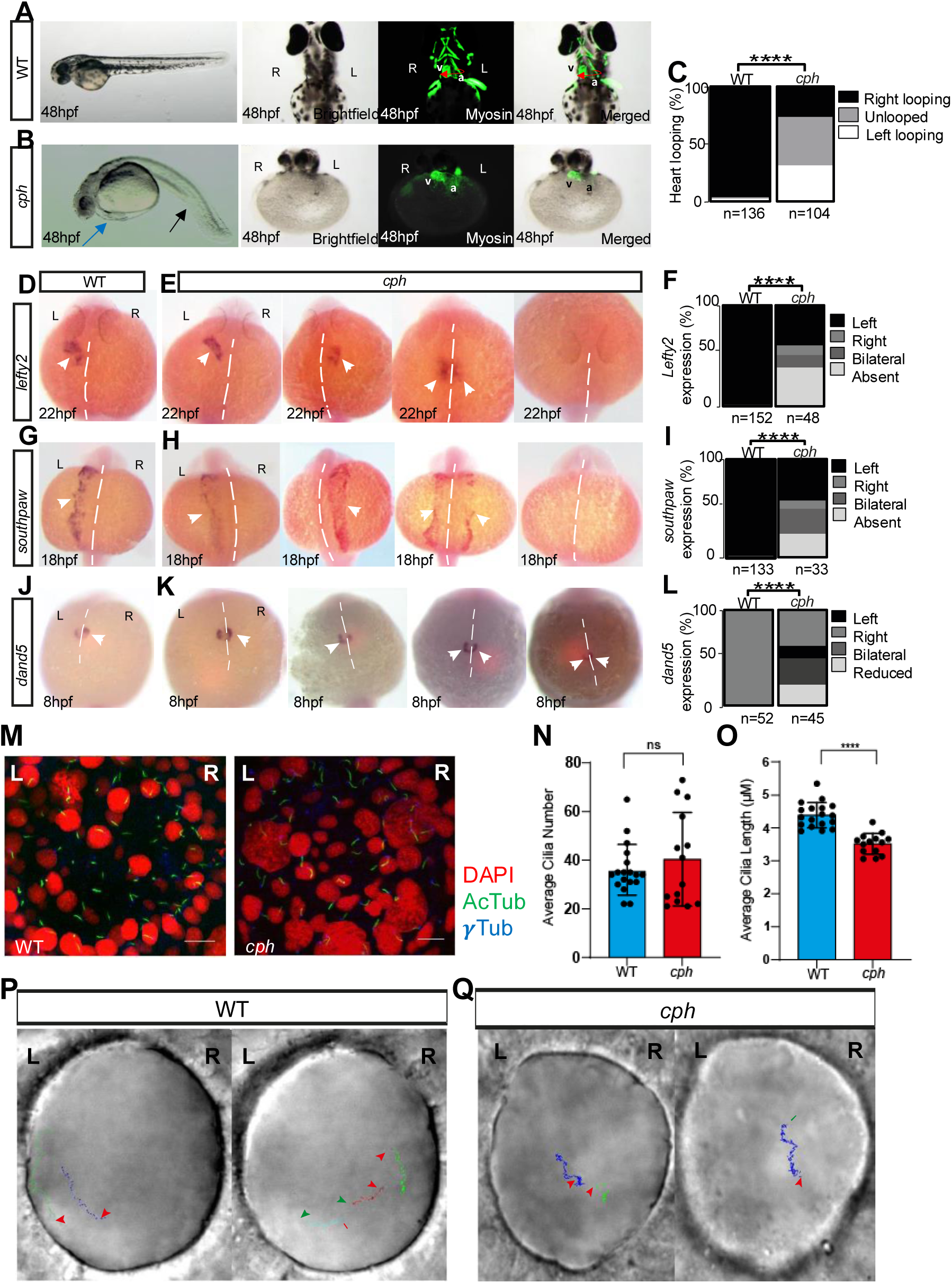
Developmental laterality and cilia defects in *cph* zebrafish. A) Wild type zebrafish at 48hpf with straight body axis (left panel). MF20 staining to identify heart tube is shown in green (right two panels). Red arrow shows looping direction of heart. B) *cph* mutant zebrafish with curled down body axis (black arrow) and pericardial oedema (blue arrow) indicative of cilia defects. MF20 staining shows delayed heart tube looping (green; right two panels). C) *cph* mutant fish show a significance increase in heart looping defects compared to wild type controls. D) Expression of the laterality gene *lefty2* in wild type zebrafish at 22hpf (white arrowheads indicate location of gene expression in panels E-L). E) Gene expression patterns of the laterality gene *lefty2* are altered in *cph* mutant fish, showing a variety of expression patterns. F) Statistical summary of *lefty2* expression pattern defects in *cph* fish. G) Expression of the laterality gene *southpaw* in wild type zebrafish at 18hpf. H) Gene expression patterns of the laterality gene *southpaw* are altered in *cph* mutant fish, showing a variety of expression patterns. I) Statistical summary of *southpaw* expression pattern defects in *cph* fish. J) Expression of the laterality gene *dand5* in wild type zebrafish at 8hpf. K) Gene expression patterns of the laterality gene *dand5* are altered in *cph* mutant fish, showing a variety of expression patterns. L) Statistical summary of *dand5* expression pattern defects in *cph* fish. M) Acetylated tubulin (green) and gamma tubulin (blue) staining in wild type fish KV (left) and *cph* fish KV (right). DAPI staining for nuclei is shown in red (Scale bar = 10µ). N) Quantification of average cilia number per KV between wild type control and *cph* fish. O) Quantification of average cilia length within individual KVs demonstrates *cph* KV cilia are significantly shorter than wild type KV cilia (n=14 and n=18 KVs respectively). Data are represented as mean ± SD. Significance was evaluated using unpaired t-test. P<0.0001 P) Still images from videos showing particle movement in wild type fish KV (representative of n=5, Scale bar = 10µm, Time lapsed = ∼3s). Q) Still images from movies showing particle movement in *cph* fish KV (representative of n=5, Scale bar = 10µm, Time lapsed = ∼3s). Green and red arrows indicate the start and end position of the bead, respectively. L left, R right.

Laterality gene expression was altered in *cph* homozygous mutant zebrafish in a manner resembling *Prpf8^N1531S^* mutant mouse embryos. Homozygous *cph* mutant embryos had absent or mis-localised expression of *lefty2* (*lft2*) (Fig. 4d-f) and *southpaw* (*spaw*), a zebrafish *Nodal* homologue (Fig. 4g-i). Additionally, *dand5,* the zebrafish *Cerl2* homologue, which is normally expressed more prominently on the right side of the embryo (Fig. 4j), displayed altered expression patterns in *cph* embryos (Fig. 4k-l). The range of abnormal expression patterns amongst mutant embryos for these key laterality genes is reminiscent of expression defects found in *Prpf8^N1531S^*mouse mutants. Overall, expression defects and heart looping abnormalities found in *cph* zebrafish suggest that the role of PRPF8 in laterality establishment and heart looping is conserved amongst vertebrates.

We next evaluated cilia formation and function in Kupffer’s vesicle (KV), the zebrafish LRO. Within KVs of wild type and *cph* zebrafish, motile cilia were present (Fig 4m), and no significant reduction in overall number of cilia was found in mutants compared to wild type controls (Fig 4n). However, there was a significant reduction in cilia length in *cph* mutants relative to wild type controls (Fig 4o; unpaired t-test. P<0.0001), similar to our findings in *Prpf8^N1531S^* mutant mouse embryos (Fig 2i). High speed live video imaging of KVs of wild type and *cph* homozygous mutants revealed that *cph* embryos exhibited a wide variation in individual cilia motility patterns. Some cilia were motile, similar to wild type (Supplemental Movie File 5), but others displayed abnormal motility, and immotile cilia were also observed (Supplemental Movie File 6). Tracking movement of endogenous particles detectable in KV fluid revealed a net anticlockwise movement in control KVs (Fig 4p, Supplemental Movie File 7), but no such net movement was observed in *cph* mutant KVs (Fig 4q, Supplemental Movie File 8), indicating a disruption in flow.

### Effect of the *Prpf8^N1531S^* mutation on mRNA splicing

Since PRPF8 is a component of the U5 snRNP that forms the spliceosomal catalytic centre, we investigated whether the *Prpf8^N1531S^* substitution affected splicing. Yeast cells containing a *N1603S* missense substitution in yeast Prp8 (homologue of PRPF8), analogous to the mouse *Prpf8^N1531S^* allele, grew similarly to wild type yeast, indicating that the Prp8^N1603S^ mutation is viable (Fig. 5a). We used the *ACT1-CUP1* splicing reporter (53) in Prp8^N1603S^ mutants to determine which step of splicing was affected. Primer extension analysis of the *ACT1-CUP1* splicing reporter revealed that with a branch site mutation (A259C), increased lariat intermediate was detected in Prp8^N1603S^ cells compared to Prp8^WT^ (Fig. 5b), indicating the first step of splicing had occurred. However, with a 3’ splice site mutation (A302U), reduced levels of mature mRNA were present in Prp8^N1603S^ cells (Fig. 5b), indicative of a defect in the second step of splicing. This assay demonstrated that the Prp8^N1603S^ mutant is a first step allele of Prp8, as it promoted the first step of splicing at the expense of the second step. We then examined growth of Prp8^N1603S^ vs Prp8^WT^ yeast colonies with *ACT1-CUP1* reporters with either 3’ splice site, 5’ splice site or branch point site mutations (Fig. 5c) on media with increasing concentrations of copper. Copper tolerance is correlated with the amount of correctly spliced mature *CUP1* transcript from the *ACT1-CUP1* construct and therefore the efficiency of splicing (53). Prp8^N1603S^ cells could not tolerate copper as well as Prp8^WT^ cells, regardless of where the mutation was in the *ACT1-CUP1* reporter construct. These data suggest that the Prpf8^N1531S^ allele does not suppress any splice site mutations and is likely to be a hypomorphic mutation.

**Fig 5.**
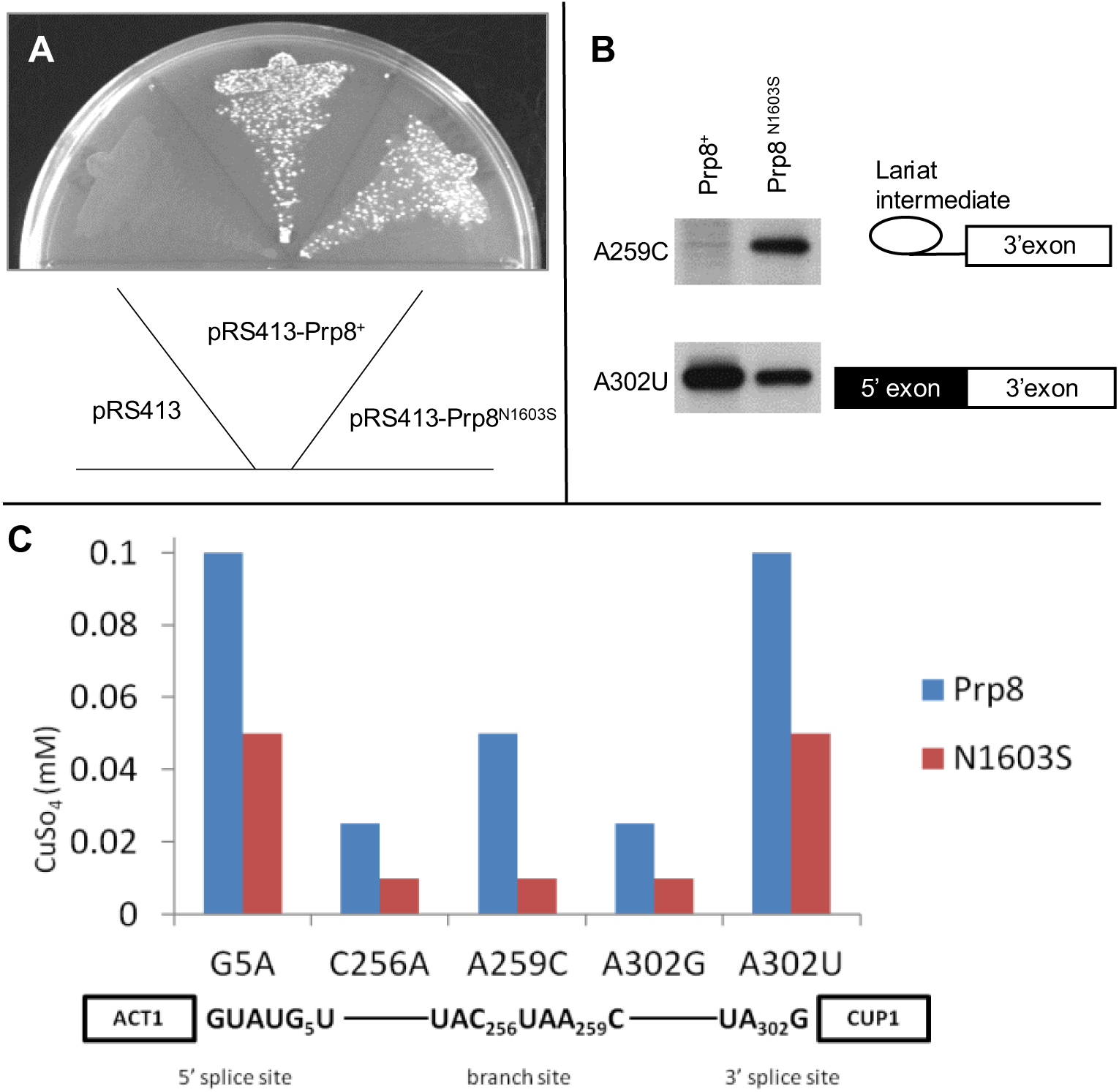
Investigation of Prp8 N1603S substitution phenotype in yeast. A) Growth of haploid yeast strains with genomic *PRP8* deletion and complementing pRS413 plasmids with either no PRP8, PRP8 wild-type (PRP8*) or PRP8 N1603S. Left sector: Strains transformed with empty pRS413 fail to grow, confirming genomic *PRP8* deletion is non-viable; middle and right sectors: yeast strains are viable when transformed with pRS413 containing either wild type or mutated (N1603S) *PRP8*. Plates were grown at 30°C. B) Primer extension analysis in Prp8 and Prp8^N1603S^ strains containing *ACT1-CUP1* reporters. Top panel displays lariat accumulation in the *ACT1-CUP1* A259C branch site reporter in the Prp8^N1603S^ strain. Bottom panel displays depletion of mature mRNA product in the *ACT1-CUP1* A302U 3’SS reporter in the Prp8^N1603S^ strain. These results are consistent with the N1603S mutation representing a first step allele of Prp8. C) Comparison of splicing efficiencies using *ACT1-CUP1* splicing reporters. More efficient splicing of a reporter correlates with increased copper resistance and therefore growth at higher concentrations of copper sulphate (CuSO_4_). Haploid yeast strains with *CUP1* and *PRP8* deletions with either wild type or N1603S Prp8 complementing plasmids were transformed with *ACT1-CUP1* splicing reporters containing mutations at the 5’SS (G5A), BPS (C256A and A259C) and the 3’SS (A302G and A302U) and grown on plates with increasing concentrations of CuSO_4_; in each case Prp8 N1603S was less efficient at splicing the reporter compared to wild type Prp8. Graphs indicate the highest concentration of copper where all three replicates survived.

### RNA-seq analysis to identify splicing defects in *Prpf8^N1531S^* mutant mouse embryos

Mutations in PRPF8 are reported to cause mis-splicing in yeast, zebrafish and humans (7, 42, 52, 54). The yeast data we present suggest that the *Prpf8^N1531S^* mutation could cause mis-splicing in mouse embryos. We therefore performed bulk RNA-seq analysis on individual whole embryos at E10.5: wild type (n=3), E10.5 *Prpf8^N1531S^* mutants with correct heart looping (n=3), and E10.5 *Prpf8^N1531S^* mutants with reversed heart looping (n=3). Analysis of differential exon skipping events between the two strains using the rMATS software package (55) identified 498 statistically significant mid-penetrance exon skipping events (average difference in exon inclusion between strains ≥ 20%; FDR-corrected p-value<0.05), across 431 unique genes (Supplemental Table 2). Gene ontology (GO) enrichment analysis revealed the most significant enrichment (24/51 genes, 2.43-fold) to be for genes associated with the biological process “plasma membrane bounded cell projection morphogenesis”, consistent with a model of ciliary dysfunction (Supplemental Table 3).

To further narrow the field of key candidate genes that could underlie the laterality defects detected in *Prpf8^N1531S^* mutant mice, individual GO terms were attached to differentially spliced genes identified by rMATS. Of the 431 unique genes harbouring exon skipping events, three - *Arl13b*, *Rfx3* and *Rpgrip1l* - were upstream of or directly involved in “determination of left/right symmetry”, although only *Arl13b* was associated with “heart looping” and “left/right axis specification”. Because *Arl13b* is involved in both general L-R axis determination and cardiac looping, and has known roles in the function of motile and non-motile cilia, the *Arl13b* exon skipping event was prioritised for additional investigation.

In *Prpf8^N1531S^* homozygous mutants, there is splicing between exons 8 and 10 of the *Arl13b* gene in approximately 51% of reads mapping to the exon 8 splice donor site, whereas in wild type embryos the corresponding event is detected in approximately 30% of reads (Fig. 6a). This exon skipping event deletes the 69bp in-frame exon 9 from the transcript. An analysis of the *Arl13b* transcript revealed that the omission of exon 9 is predicted to remove a 23 amino acid segment of the Proline Rich Region from the C-terminal end of ARL13B (Fig. 6b). We have termed this *Arl13b* transcript, lacking exon 9, the “short isoform”, and the transcript including exon 9 the “long isoform”. The short isoform is not annotated in the fish or mouse UCSC genome browsers, although a single mouse EST clone (DT914850) does have the same sequence as the *Arl13b* short isoform we have identified. To our knowledge the function of this isoform has not been studied previously. Using a single pair of PCR primers located in exons 8 and 10, we amplified both short and long *Arl13b* isoforms by RT-PCR in wildtype mouse and zebrafish embryos (Fig. 6c). There was an increased ratio of the *Arl13b* short isoform transcript to long transcript in both *Prpf8^N1531S^*homozygous mouse and *cph* homozygous mutant fish embryos (Fig. 6c). Sequencing of these RT-PCR products revealed the precise deletion of exon 9 from the short isoform product (Fig. 6d). We also detected the *Arl13b* short isoform in multiple adult mouse tissues, most notably in the testes (Fig. 6e-f).

**Fig 6.**
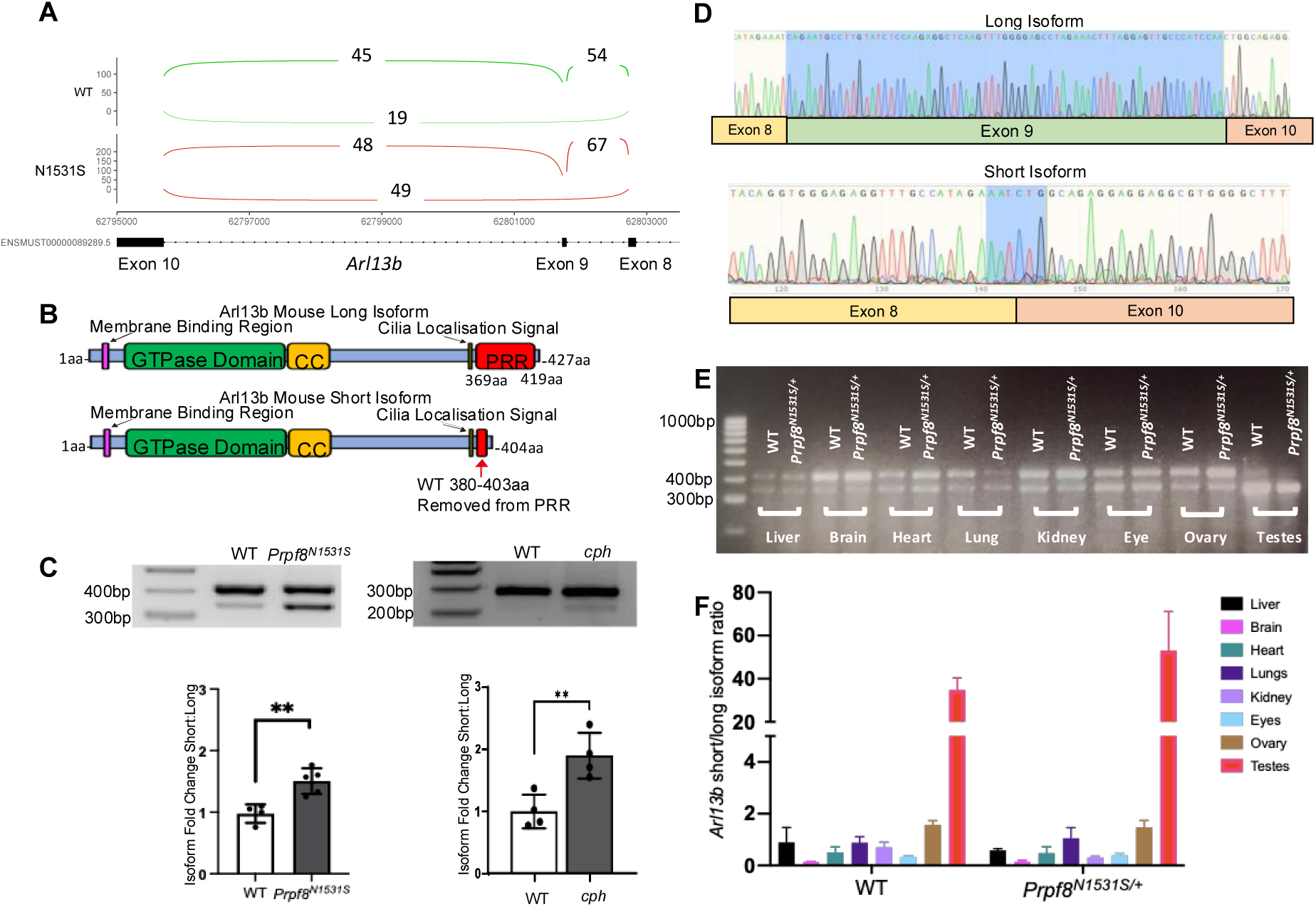
*Arl13b* transcript alterations present in *Prpf8* mutant mice and fish. A) Sashimi plot illustrating relative read coverage of splice junctions between exons 8 and 10 of the murine *Arl13b* transcript in three WT mice (top, green) and three *Prpf8^N1531S^* homozygous mouse mutants (bottom, red). Numbers represent the mean read coverage across each junction for the three replicates of each genotype. *Prpf8^N1531S^* homozygous mutants exhibit a marked increase in skipping of exon 9, producing greater levels of the *Arl13b* short isoform. B) Schematic representation of ARL13B predicted protein products found in wild type embryos and *Prpf8^N1531S^*mouse mutants. The region of the transcript encoding amino acids 380-403 from the wild type sequence is removed from the short isoform. C) RT-PCR using a single primer pair displaying long and short isoforms of *Arl13b* in mouse (left) and zebrafish (right) embryos. Quantification of the relative proportions of the *Arl13b* short and long isoforms plotted as fold change relative to wild type are shown below the agarose gels. D) Sequencing chromatograph of *arl13b* RT-PCR product from wild type zebrafish (top) and *cph* embryos (bottom). The region corresponding to exon 9 is highlighted in the long isoform sequence. The junction between exon 8 and 10 is highlighted in the short isoform sequence. E) RT-PCR using a single primer pair displaying *Arl13b* long and short isoform distribution in wild type and *Prpf8^N1531S^*heterozygous adult mouse organs. F) Quantification of the ratio of relative expression levels of *Arl13b* long and short isoform in wild type and *Prpf8^N1531S^* heterozygous adult mouse organs.

### The ARL13B short isoform localises to cilia

To determine if the shift in *ARL13B* isoform usage could affect phenotype, we generated *ARL13B* short or long isoform human transcript sequences with N-terminal GFP fusions. These were transfected into hTERT-RPE1 cells for comparison with endogenous ARL13B. Endogenous ARL13B co-localised with acetylated tubulin at the cilium in hTERT-RPE1 cells (Fig. 7a left). We detected protein produced from the GFP-tagged ARL13B short isoform co-localised with endogenous acetylated tubulin at the cilium in hTERT-RPE1 cells (Fig. 7a centre), demonstrating the short isoform produces a stable protein that is localised correctly. Similar localisation results were seen for the GFP tagged ARL13B long isoform (Fig. 7a right). The short isoform cilia localisation is consistent with the ARL13B cilia localisation signal being within amino acids 347-363 (56), which are included in the short isoform transcript.

**Fig 7.**
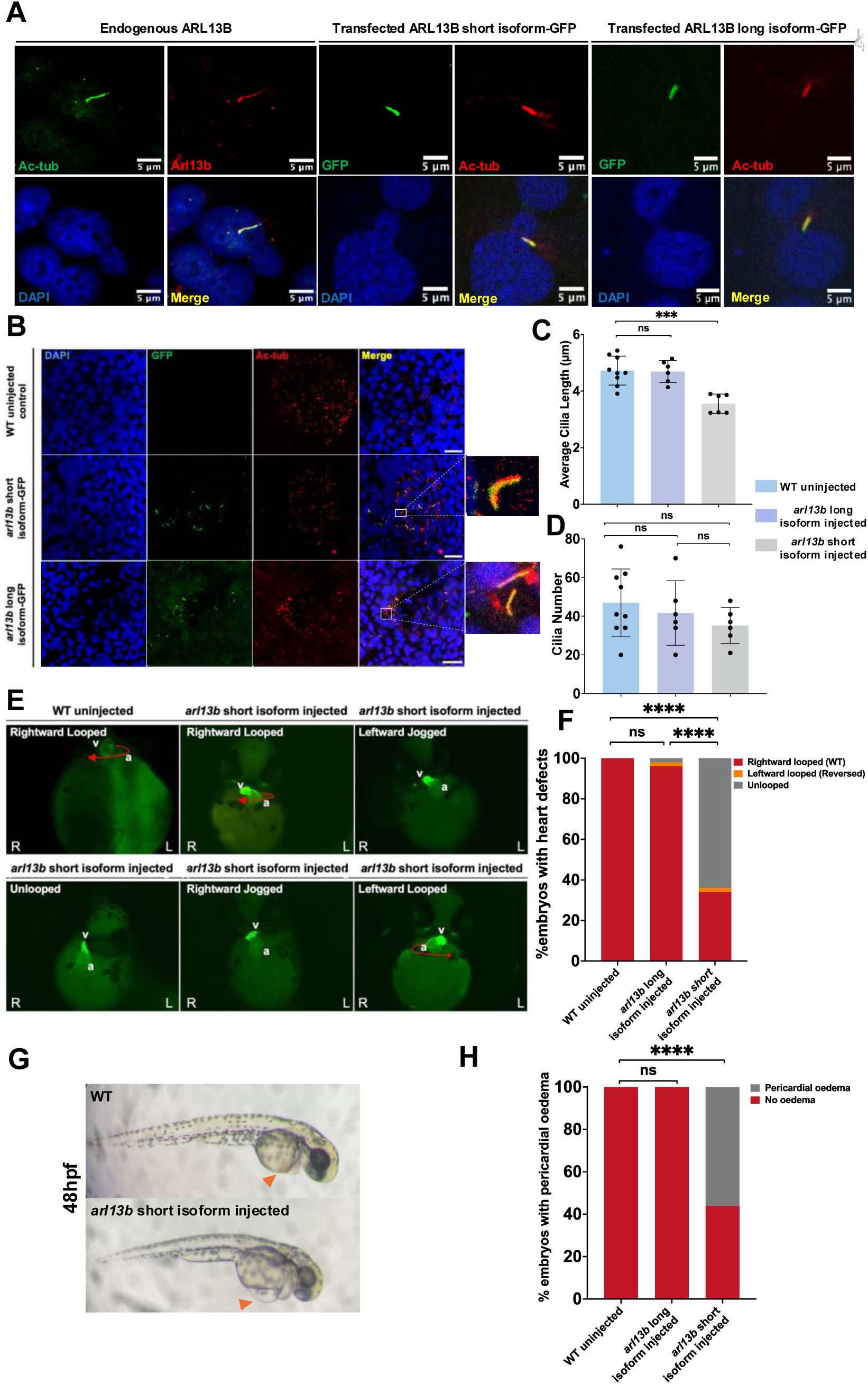
Analysis of *ARL13B* short and long isoform over-expression in human cells and wild type zebrafish embryos. A) Co-localization of human acetylated tubulin (green) and endogenous ARL13B (red; left panels); co-localization of transfected GFP-tagged human ARL13B short isoform (green) with acetylated tubulin (red; centre panels); co-localisation of transfected GFP-tagged human ARL13B long isoform (green) with acetylated tubulin (red; right panels) in hTERT-RPE1 cells. Nuclei are stained with DAPI (blue). Scale bar 5μm. B) Immunofluorescence images of KV cilia in three groups: wild type uninjected embryos (top), embryos injected with *arl13b*-*gfp* short isoform mRNA (middle), and embryos injected with *arl13b-gfp* long isoform mRNA (bottom). Cilia are stained with acetylated tubulin (labelling axonemes), and GFP to label the injected Arl13b protein isoforms. Nuclei are stained with DAPI. C) Quantification of average cilia length per KV in wild type uninjected control embryos (n=9); *arl13b* long isoform injected embryos (n=6), and *arl13b* short isoform injected embryos (n=6). No significant difference was detected between the *arl13b* long isoform injected embryos and the uninjected wildtype embryos (one-way ANOVA, p > 0.05). Cilia length in *arl13b* short isoform injected embryos was significantly decreased compared to uninjected wildtype embryos (one-way ANOVA, p <0.05). Scale bar = 20μm. D) Quantification of cilia number per KV in control and injected embryos. No significant differences were detected between groups (one-way ANOVA p >0.05). E) Heart directionality at 48hpf detected by MF20 immunofluorescence (red arrows). Some *arl13b* short isoform injected injected embryos show delayed cardiac development and have not achieved fully looped hearts at 48hpf, but resemble the jogging asymmetry typically present at 24hpf. Injection condition is shown above the images, observed heart directionality is shown on the images. F) A significant difference in heart defects frequency was presented among the three group (Chi-square test, p< 0.0001). Pairwise comparisons then were performed between groups. Embryos injected with the *arl13b* short isoform mRNA (n=50) displayed a significantly increased percentage of heart directionality defects compared with the *arl13b* long isoform mRNA injected (n=50) or wild type uninjected group (n=50) (Fisher’s exact test, p < 0.0001), while there is no significant difference of heart defect frequency observed between the *arl13b* long isoform mRNA injected group and the wildtype uninjected group (Fisher’s exact test, p = 0.4949). G) Brightfield image of wild type uninjected embryo (bottom) at 48hpf lacking oedema in pericardium (orange arrowhead). *arl13b* short isoform mRNA injected embryo (bottom) at 48hpf shows pericardial oedema (orange arrowhead). H) Wild type embryos (n=50) do not display pericardial oedema at 48hpf, nor do embryos injected with the *arl13b* long isoform (n=50). Embryos injected with the *arl13b* short isoform mRNA (n=50) showed a significant increase in pericardial oedema compared with the uninjected control group (Fisher’s exact test, p < 0.0001).

### Over-expression of the *arl13b* short isoform in zebrafish embryos mimics the *cph* mutant phenotype

We next sought to determine if increased expression of the zebrafish *arl13b* short isoform causes cilia defects. We injected wild type zebrafish embryos with mRNA encoding a C-terminal Arl13b-GFP fusion protein of either the short or long isoform. At 12 hours post-injection, embryos exhibiting GFP were selected for further experiments. Half of the selected embryos were fixed immediately for immunofluorescence to analyse Arl13b protein localisation and KV cilia length, while the remaining embryos were monitored until 48 hours post-injection to investigate phenotypes indicative of cilia dysfunction: cardiac looping and pericardial oedema. In injected zebrafish embryos both Arl13b long and short isoform GFP fusion proteins localised to KV cilia, showing co-localisation with acetylated tubulin (Fig 7b). Measuring KV cilia length for each injection group, and plotting the average cilia length per KV, revealed no significant change in KV cilia length in wild type fish injected with 80ng/ul of *arl13b* long isoform GFP-tagged mRNA as compared to uninjected control fish (Fig 7c; One-way ANOVA test, p=0.9901). However, cilia within the KV of fish injected with the *arl13b* short isoform were significantly shorter than those in the uninjected control group (Fig 7c; One-way ANOVA test, p=0.0002), similar to findings for *cph* KV cilia length. The number of cilia per KV was not significantly different between any of the groups (Fig 7d).

We next analysed cardiac looping at 48hpf in embryos injected with *arl13b* long and short isoforms compared to uninjected controls. Immunofluorescence microscopy for the cardiac marker MF20 revealed varied heart looping directionality. The uninjected control group heart tube was correctly looped (Fig 7e top left); embryos from *arl13b* short isoform mRNA injection showed a variety of cardiac laterality patterns including correct looping (34%), delayed development and incomplete looping (66%) and reversed looping (2%) (Fig 7e; heart looping direction listed on top left of image). No significant changes were seen in *arl13b* long isoform injected embryos (Fig 7f; Fisher’s exact test, p= 0.4949), whereas a significant proportion of *arl13b* short isoform injected embryos displayed cardiac looping abnormalities (Fig 7f; Fisher’s exact test p<0.0001). Bright-field images of wild type uninjected fish captured at 48hpf showed the expected cardiac morphology (Fig 7g top), whereas 56% of *arl13b* short isoform injected embryos displayed pericardial oedema (Fig 7g bottom). The percentage of the *arl13b* short isoform injected embryos showing pericardial oedema at 48hpf was significantly increased compared to uninjected controls (Fig 7h; Fisher’s exact test, p<0.0001).

### Over-expression of *arl13b* long isoform rescued phenotypic defects of *prpf8* loss in zebrafish embryos

To determine if disruption of *arl13b* isoform usage is, at least in part, the mechanism underlying the cardiac laterality defects in *cph* mutant fish, we utilised a *prpf8* splice blocking morpholino (sbMO) targeting exon 7 (Supplemental Fig. 3), which provokes cardiac laterality defects in injected zebrafish (57). KV cilia were identified by acetylated tubulin immunofluorescence (Fig.8a). Compared to WT uninjected control embryos, the sbMO injected embryo had significantly decreased KV cilia length (Fig. 8b; Unpaired t-test, P<0.0001), but no significant difference was detected in cilia number per KV (Fig. 8c; Unpaired t-test; p = 0.4868). Immunofluorescence microscopy at 48hpf showed cardiac looping defects in embryos injected with *prpf8* sbMO (Fig. 8d), while co-injection of *prpf8* sbMO with *arl13b* long isoform mRNA reduced the frequency of cardiac looping defects. Co-injection of *prpf8* sbMO with *arl13b* short isoform mRNA did not rescue cardiac development. Injection of a control MO showed no cardiac defects (Fig. 8d). Quantification of cardiac defects revealed significant differences between the sbMO and control embryos (Fig. 8e, Fisher’s exact test p <0.0001), but not between embryos co-injected with *prpf8* sbMO and *arl13b* long isoform mRNA when compared to control embryos (Fig. 8e; Fisher’s exact test, p >0.05). Additionally, a high incidence of severe pericardial oedema and tail curvature defects was seen in embryos injected solely with *prpf8* sbMO, but embryos co-injected with *prpf8* sbMO and *arl13b* long isoform mRNA exhibited a significant reduction in the percentage of these defects (Fig. 8f-g; Fisher’s exact test p <0.0001). The 72hpf survival rate for *prpf8* sbMO embryos co-injected with *arl13b* long isoform mRNA was significantly increased compared to *prpf8* sbMO injection alone (Fig. 8h; Fisher’s exact test p <0.0001). Morphological analysis of wild type uninjected control embryos or embryos co-injected with *prpf8* sbMO and *arl13b* long isoform mRNA did not reveal obvious anatomical defects (Fig. 8i) at 72hpf, whereas *prpf8* sbMO injected embryos exhibited pericardial oedema (black arrowhead), head morphological defects (blue arrowhead), eye defects (green arrowhead) and tail defects (orange arrowhead), as did embryos co-injected with *prpf8* sbMO and *arl13b* short isoform mRNA.

**Fig 8.**
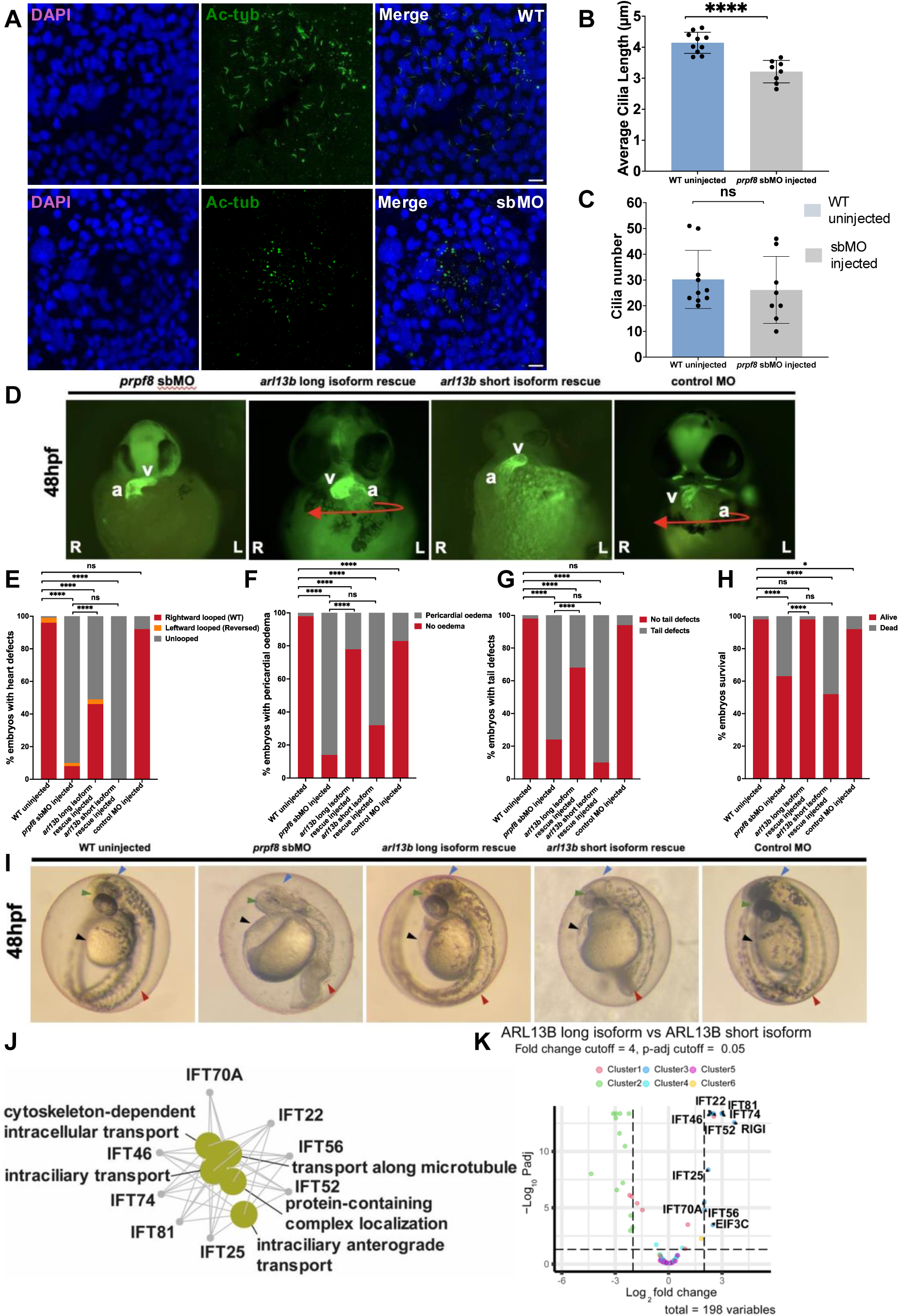
Analysis of *Prpf8* sbMO knockdown phenotypes and rescue by *arl13B* long isoform mRNA co-injection. A) Immunofluorescence of KV cilia stained using an antibody against acetylated tubulin labelling axonemes (green) in WT and *prpf8* sbMO knockdown embryos. Nuclei are stained with DAPI (blue). Scale bar is 10μm. B) Quantification of cilia length per KV in WT (n=10) and *prpf8* splice blocking morpholino (sbMO) knockdown embryos (n=8). All cilia detected with acetylated tubulin staining were measured, and the average length per KV is plotted. *prpf8* sbMO knockdown embryos show a significant decrease in cilia length (Unpaired t-test, p < 0.0001). C) Quantification of cilia number per KV showed no significant difference between groups (Unpaired t-test; p =0.4868). D) Cardiac looping morphogenesis detected by MF20 immunofluorescence. sbMO embryos and *arl13b* short isoform mRNA rescue embryos showed heart looping defects. The co-injection of *arl13b* long isoform mRNA with sbMO partially rescued heart looping defects. Control MO injected embryos did not display heart directionality defects. E-H) Analysis of phenotypes in wildtype uninjected embryos (n=94), *prpf8* sbMO knockdown embryos (n=104), sbMO co-injection with *arl13b* short isoform mRNA (n=44), sbMO co-injection with *arl13b* long isoform mRNA (n=79), and control MO injected embryos (n=84). Statistical analysis was first performed using a Chi-square test to assess morphological defects among five groups. Where significant difference was observed, pairwise comparisons were conducted between two groups by Fisher’s test. E) The sbMO knockdown embryos with *arl13b* long isoform mRNA injection displayed a significantly decreased percentage of heart directionality defects compared to the sbMO knockdown group (Fisher’s exact test, p < 0.0001). F) The percentage of pericardial oedema at 48hpf is significantly decreased in the *arl13b* long isoform mRNA injection rescue group compared to the sbMO knockdown group (Fisher’s exact test, p < 0.0001). G) The percentage of tail defects is significantly decreased in the *arl13b* long isoform overexpression rescue group compared to the sbMO knockdown group (Fisher’s exact test, p < 0.0001). H) Embryo survival at 72 hpf is significantly increased in the long isoform co-injection group compared to the sbMO knockdown group (Fisher’s exact test, p < 0.0001). I) Brightfield images of WT embryo, *prpf8* sbMO, *arl13b* long isoform co-injected embryo, *arl13b* short isoform co-injected embryo and control MO injected embryo. Black arrowheads indicate pericardial oedema, blue arrowheads indicate head morphological defects, green arrowheads indicate eye defects, and red arrowheads indicate tail defects seen in *prpf8* sbMO injected embryos. J) Network analysis of enriched biological processes at GO Level 5 for proteins binding to the long isoform of ARL13B. Many processes related to cilia function are present. K) Volcano plot of mass spectrometry data showing proteins that are enriched for binding the long isoform of ARL13B compared to the short isoform, based on a fold-change cutoff of 4 and an adjusted p-value cutoff of 0.05. The X-axis represents the log2 fold change, and the Y-axis represents the log2 adjusted p-value. Note that many members of the IFT-B complex are significantly enriched in this comparison.

### The ARL13B short isoform displays reduced binding to IFT-B1 complex proteins

The short isoform of ARL13B lacks part of the proline rich region near the C-terminus of the protein. The proline rich region binds IFT46 and IFT56 (58), components of the IFT-B1 complex involved in anterograde ciliary trafficking (59). We therefore sought to examine differences in protein interactions between the long and short isoforms of ARL13B. Using a GFP-trap system (60), we expressed either the GFP tag alone, or human long or short ARL13B-GFP fusion protein isoforms in RPE1 cells, followed by GFP affinity purification-mass spectrometry of three biological replicates. Network analysis of biological processes that are enriched in proteins associated with the long isoform compared to the short isoform revealed a network associated with cilia function (Fig. 8j; Supplemental Fig. 5). The short isoform of ARL13B is depleted for binding to IFT22, IFT25, IFT46, IFT52, IFT56, IFT70a, IFT74, and IFT81, which are all proteins found within the IFT-B1 complex (Fig. 8k; Supplemental Fig. 5). Overall, these data suggest that the correct balance between ARL13B isoforms is critical for maintaining cilia structure and function during development.

## Discussion

We report a novel, evolutionarily conserved role for *Prpf8* in LRO cilia function and determination of organ laterality in vertebrate embryos. We identified a shift in *Arl13b* isoform usage as a potential mechanism explaining the laterality defects observed in *Prpf8* mutant mouse and zebrafish embryos. Consistent with this finding, *Arl13b* mouse mutants (61, 62) display cardiac looping defects, highly reminiscent of *Prpf8^N1531S^* embryos described here (Fig 1), supporting the hypothesis that altered *Arl13b* isoform usage can perturb left-right axis formation and embryonic development. We did not observe changes in PRPF8 protein localisation to cilia (Supplemental Fig 2), suggesting that any as yet undefined ciliary, non-splicing roles of PRPF8 (12) may be intact in the *Prpf8^N1531S^* homozygous mouse mutant.

To our knowledge, other characterised mouse strains with mutations in *Prpf8* do not resemble the *Prpf8^N153S^* embryonic lethal laterality defect phenotype that we have detected (63–65). Previously characterised mutants were generated to model human RP, containing variants in the globular Jab1/MPN domain of PRPF8 (10). The N1531S mutation lies within the linker domain of PRPF8, near the endonuclease-like domain (43), and thus may have a different effect on PRPF8 function than mutations in the Jab1/MPN domain. Although many missense variants throughout yeast Prp8 have been characterised (42), additional study is needed in mammalian models to reveal the complexity of transcriptome and phenotypic defects that may arise from variants in differing protein domains. However, an iPSC model of a heterozygous *PRPF8* c.6926 A > C (p.H2309P) RP-associated mutation caused impaired alternative splicing and enhanced cryptic splicing predominantly in ciliary and retinal-specific transcripts (11), which is consistent with findings from the mouse N1531S variant. Although we have shown several aspects of the *Prpf8^N1531S^* and *cph* mutant phenotypes are recapitulated by over-expression of the *arl13b* short isoform mRNA, we cannot exclude contributions from other genes to the mutant phenotypes. For example, we detected splicing alterations in the transcription factor *Rfx3* in *Prpf8^N1531S^* mutants. *Rfx3* knockout embryos have laterality and node cilia elongation defects (66), but their overall morphology, frequency of situs inversus and incompletely penetrant embryonic lethality (66) are dissimilar to the phenotypes of *Prpf8^N1531S^* and *cph* mutants.

We demonstrated that increased usage of a previously uncharacterised transcript isoform of *arl13b*, lacking exon 9, disrupted ciliogenesis and provoked laterality defects when over-expressed in wild type zebrafish. We detect the short isoform in all tissues tested, in varying ratios compared to the long isoform. Additional research is needed to reveal the roles of the two different ARL13B isoforms in different tissues and cell types. In humans, pathological variants in many ciliary genes, including *ARL13B*, cause Joubert syndrome (6, 67, 68), a well-recognized ciliopathy (69). Laterality defects have been reported in some Joubert syndrome patients, although individuals with laterality defects did not have variants in *ARL13B* (70). Interactions between ARL13B and the IFT complex govern the speed of IFT entry into cilia, and consequently, ciliary length (36, 71). Mutations in *Arl13b* and some IFT components in zebrafish and mouse have been noted to result in short cilia (62, 72–74), suggesting that a similar mechanism may play a role in the reduction in cilia length in zebrafish KV cilia in embryos over-expressing the *arl13B* short isoform mRNA. Additionally, missense variants in IFT52 cause cilia elongation defects in IMCD3 cells (75). Although the short isoform of the ARL13B protein identified in *Prpf8^N1531S^* and *cph* mutants localises to cilia, we demonstrated that its interactions with IFT-B components, including IFT52, are reduced. Thus, the shorter cilia detected in mutant LROs are consistent with a loss of IFT function.

Cilia motility is driven by dynein function. We found motility defects in the LRO cilia in both *Prpf8^N1531S^* mutant mouse and *cph* fish embryos, yet the mouse node cilia appear to have intact dynein inner and outer arms. Defective cilia motility with preservation of dynein protein cilia localisation has previously been reported for *Dnaaf1^m4Bei^* and *Lrrc48^m6Bei^* mouse mutants (76). Therefore, although the mechanism for cilia motility defects in *Prpf8* mutants is not fully understood, there is a precedent for the disruption of cilia motility without dynein arrangement defects. Additionally, the shift in ARL13B isoform usage results in reduced binding between ARL13B and IFT-B complex proteins, including IFT74, IFT22 and IFT81. IFT74 has physical interactions with IFT22 and IFT81, and a binary interaction with the inner dynein arm protein DNALI1 (77) proposed to be required for cilia motility (78). Patients with hypomorphic IFT74 variants show cilia motility defects, although not laterality defects (79). A mouse IFT74 mouse null mutant shows mid-gestation lethality and cardiac oedema, but laterality defects were not reported for these mutants (80). To our knowledge, it is not clear if these proteins are localised to the LRO cilia during development, but this is an area for future investigation. There is high conservation of protein content between cilia of different types and different localisations (81), which notably includes ARL13B and the IFT proteins found to be affected in PRPF8 mutants. Together, our findings suggest a mechanistic link between altered PRPF8 protein function in the spliceosome, aberrant splicing of *Arl13b* transcript and abnormalities in IFT and cilia function, culminating in conserved laterality defects in mutant embryos.

## Materials and Methods

### *l11Jus27* mouse model

The *Prpf8^N1531S^* strain is derived from the *l11Jus27* strain (32) and was backcrossed for at least 2 generations to the 129S5/SvEvBrd line carrying the chromosomal inversion Inv(11)8Brd*^Trp-Wnt3^*, and then maintained through sibling intercrosses for at least 6 generations. Mouse colonies were maintained at the Biological Service Facility at the University of Manchester, UK, according to Home Office requirements and with local ethical approval (project licence number PP3720525). Mice were euthanised using a Schedule 1 method following UK Home Office regulations. Genotyping was performed as described (41).

### Mouse embryo morphological analysis

Embryos were isolated from timed pregnant female mice by dissection in PBS. Morphology was analysed using a Leica dissecting microscope. Whole embryos were incubated in 1 ng/mL propidium iodide staining solution for 5 min, followed by 3 × 5 minute washes in PBS, before imaging with a DAPI filter to identify heart tube laterality.

### Genome sequencing

Genomic DNA was extracted from an E10.5 *l11Jus27* homozygous embryo using ISOLATE II Genomic DNA Kit (Bioline). 1 μg of DNA was sequenced at The Manchester Centre for Genomic Medicine (MCGM), Saint Mary’s Hospital, Manchester, UK, on the Illumina HiSeq platform. Reads were then aligned to the mouse MGSCv37 (mm9) genome assembly. Variants in the candidate region on chr 11 were analysed. Known variants in 129 and B6 mouse strains were excluded, as well as variants present in fewer than 50% of reads. The remaining variants were annotated with Ensembl Variant Effect Predictor (VEP). A single novel exonic mutation was present in 100% of mapped reads: an A to G transition at position Chr11: 75,391,978 (mm39). Sanger sequencing was performed to verify the sequence change in three additional individual embryos (University of Manchester Genomic Technologies Core Facility).

### Mutation Analysis

Conservation of protein sequence across different species was evaluated using multiple sequence alignments in Clustal Omega (https://www.ebi.ac.uk/Tools/msa/clustalo/). Mutation pathogenicity was predicted using SIFT (45) and PolyPhen2 (44). The Prpf8 protein Alphafold structure was downloaded from AlphFold Protein Structure Database: https://alphafold.ebi.ac.uk/entry/Q6P2Q9; and opened as entry AF-Q99PV0-F1-v4 in PyMOL software (Version 3.1.1).

### Quantitative PCR

*Prpf8* expression was quantified using a *Prpf8* TaqMan probe (Applied Biosystems Cat: 4331182) on a DNA Engine Opticon 2 continuous fluorescence detector (BioRad). The raw data were processed on Opticon monitor 3 (BioRad) software and subsequently analysed using Excel (Microsoft). Samples were normalised against expression levels of *Gapdh* (primers listed in Supplemental Table 4).

### Scanning Electron Microscopy

Embryos at were dissected in PBS and fixed in 2.5% glutaraldehyde, 4% paraformaldehyde in 0.1M HEPES (pH7.4) at 4°C from overnight up to 1 month. Embryos were then washed 5x 5 minutes in dH_2_O, then incubated for 1 hour in OsO_4_ in 0.1M HEPES (pH7.4). They were then washed 5x 5 minutes in dH_2_O and taken through an ethanol dehydration series before critical point drying. Embryos were mounted ventral side up on stubs using graphite sticky tape and conductive epoxy. Samples were imaged at high vacuum at 10KV on a Quanta FEG 250 electron microscope (FEI).

### Transmission Electron Microscopy

Embryos were fixed and processed as for SEM. Embedding, sectioning and imaging was performed as previously described (82). Samples were imaged on a Tecnai 12 Biotwin transmission electron microscope.

### Nodal cilia videomicroscopy

E8.25 embryos were dissected individually in warmed DMEM modified with HEPES without glutamine and sodium pyruvate (Sigma D6171), supplemented with 10% heat inactivated FBS (Gibco 10500064). Embryos at the 1-3 somite stage were selected for imaging and were transferred to a homemade chamber slide with the ventral node facing the microscope objective. To visualise the nodal flow, the media was replaced with a 1:100 dilution of latex microbeads (Sigma L1398) in media. A coverslip was then placed over the chamber and the slides were incubated at 37°C for 10 minutes before imaging on a Leica DMRB at 40X using a Leica MC170HD camera. Five-minute videos were taken using VGA2USB (Epiphan Video) with an MPEG4 decompressor at maximum quality. Video analysis was performed using the Manual Tracker plugin from Fiji (83). Movies were converted to .tiff stacks of 150 frames from the first minute of filming and the paths of 5 beads were tracked.

For visualising nodal cilia movement directly, embryos were imaged on a Nikon TE2000 PFS microscope using a 100x/1.49 Apo TIRF objective. Videos lasting 20 seconds were collected on a FASTCAM SA3 (Photron) at 125 frames per seconds and analysed using Fiji. Imaging data were enhanced for visual analysis by performing 11-frame running-average background subtraction followed by inversion and thresholding, using custom routines coded in MATLAB R2015a with Image Processing toolbox (Mathworks, Natick, MA). Motile cilia were counted manually by inspection of 125 frame (1 s) sequences taken at each focal plane.

### Mouse and zebrafish embryo whole mount *in situ* hybridisation

Whole mount *in situ* hybridisation was performed according to established protocols in mouse (84) and zebrafish (85). Control and mutant embryos were pooled and hybridised with digoxigenin labelled probes and incubated with alkaline phosphatase conjugated anti-digoxigenin antibodies. Hybridised embryos were then developed with NBT-BCIP until desired staining was achieved. *In situ* probes were synthesised using either plasmid or PCR templates. *In situ* probe primer sequences for PCR templates are listed in Supplemental Table 4.

### Mouse embryo immunofluorescence staining

Antibodies used for immunostaining are found in Supplemental Table 5. Embryos previously stored at −20C in methanol were rehydrated through a methanol series, washed 3x 20 minutes in PBS and blocked for 90 minutes at room temperature in 5% horse serum and 0.5% Triton X-100 in PBS. Control and mutant embryos were pooled and then incubated overnight at 4°C in 5% horse serum, 0.5% Triton X-100 in PBS with primary antibodies. Embryos were then washed in 3×20 minutes in PBS and then incubated with secondary antibodies in PBS with 1% horse serum for 2 hours at room temperature. Embryos were washed 3×10 minutes in PBS and then incubated with Cy5 labelled streptavidin in PBS for 30 minutes. Embryos were washed 3×20 minutes in PBS with 1% horse serum and then stained with DAPI in PBS, before a final 10 minute wash in ice-cold PBS. Embryos were then individually placed in dishes of PBS and the anterior portion of the embryo was removed. The posterior portion, including the node, was transferred to a homemade chamber slide, excess PBS removed, and embryos mounted in Prolong Gold Antifade (Life Sciences, P36934). The anterior portion was used to re-genotype the samples. The embryo was arranged ventral side up and a coverslip was placed on top, with gentle pressure applied to remove bubbles and excess mounting medium. Slides were cured for at least 2 days at room temperature.

Embryos were imaged on a Leica TCS SP5 AOBS upright confocal microscope using a 63x/1.40 HCX PL Apo objective and a 2.0x digital zoom using sequential imaging in a 2048×2048 format.

### Mouse Embryo Western Blot

E10.5 mouse embryos were lysed in RIPA buffer supplemented with protease inhibitors. As described earlier, protein lysates were separated by SDS-PAGE and transferred to PVDF membranes (86). Western blotting was performed using antibodies against PRPF8 and GAPDH (loading control). Signal detection was carried out using LI-COR Odyssey system and the image was quantified using the LI-COR Image Studio software.

### PRPF8 FLAG-tagged plasmid generation and transfection

IMAGE clone 5587081 containing human PRPF8 was used for generating PRPF8 constructs. Mouse and human PRPF8 proteins differ by only 3 of the total 2335 amino acids. We generated the N1531S variant by site directed mutagenesis (Q5 Site Directed Mutagenesis kit; NEB), and corrected a A759G missense sequence change present in the IMAGE clone. Primer sequences are listed in Supplemental Table 4. To correct the A759G sequence change, primers creating a new restriction site for Eco47III were utilised. Rather than creating the exact *l11Jus27* mutation (aat to agt), we used an alternate codon sequence which generated the same amino acid mutation (aat to agc), but created a BseYI restriction site allowing screening for colonies containing the correct mutation prior to sequence verification. Oligonucleotides containing a FLAG tag sequence, a start codon, a Kozak sequence and a Cla1 restriction site were cloned into the corrected wild type and the N1531S mutant PRPF8 plasmids at an AgeI restriction site.

### PRPF8 protein stability analysis

HEK293T cells were cultured as described previously (86). The cells were seeded at a density of 0.1×10^6^/well in 6 well plates for transfection. The next day the cells were transfected with 2µg of *PRPF8* WT-Flag or N1531S-Flag plasmid using Fugene HD reagent (Promega). 48hours later the transfection media was removed, and cells were treated with 100µM cycloheximide in cell culture media. The cells were lysed at different time points (0, 4, 8, 16 hours) in RIPA buffer. The immunoblotting was performed using anti Flag and anti β-Actin (ACTB) antibodies as described previously (86). Signal detection was carried out using LI-COR Odyssey system. Image was quantified using the LI-COR Image Studio software.

### Zebrafish *cph* mutants

Fish were maintained at the Institute of Molecular and Cell Biology, A*STAR Zebrafish Facility and the University of Manchester Biological Services Facility (IMCB protocol number: 221702 and Manchester project licence PP8367714) at a temperature of 28.5°C and a standard light/dark cycle of 14 hours and 10 hours respectively. Pair mating was used to obtain embryos. Embryos were incubated at 28.5°C in egg water containing methylene blue until the desired stage.

### Genotyping *cph* mutants

*cph* mutant zebrafish were genotyped by PCR amplification of exon 8 and 9 of the *prpf8* gene (52), followed by purification using QIAquick PCR and Gel Purification Kit (Cat. No 28104), and a restriction digest using AccI enzyme performed as per manufacturer’s instructions. Samples were ran on a 2% agarose gel. WT fish produced one band, heterozygotes produced 3 bands and *cph* homozygotes produced 2 bands (Supplemental Figure 6).

### Zebrafish whole mount immunofluorescence

Embryos were transferred into a glass cavity dish and rehydrated from methanol by washing in consecutively lower methanol concentrations (75%, 50%, 25%) in PBS. Embryos were subsequently washed in PBS for 3x 5 minutes. PBS was removed and embryos treated with −20°C acetone and placed at −20°C for 7 minutes (this step was not used for embryos fixed with Dent’s fix). Embryos were washed for 3x 5 minutes in room temperature PBS. Embryos were incubated for 1 hour at room temperature in blocking buffer (2% sheep serum in PBDT) on an orbital shaker at 70 rpm. Blocking buffer was removed and primary antibody diluted in PBDT added (Supplemental Table 5). Embryos were placed on an orbital shaker at 70rpm overnight at 4°C. The following morning primary antibody was removed and embryos washed in room temperature PBDT 3x 30 minutes on an orbital shaker at 70rpm. PBDT was removed and secondary antibodies diluted in fresh PBDT were added. Embryos were placed on an orbital shaker at 70rpm for 5 hours at room temperature. During the last 30 minutes DAPI was added (1:2000). Secondary antibodies were removed and embryos washed 3x 30 minutes with room temperature PBDT. PBDT was removed and embryos were submerged in 70% glycerol for storage at 4°C. Primary antibodies for immunostaining found in Supplemental Table 5.

### Measurement of Node and KV LRO Cilia

To detect nodal cilia, mouse 1-4 somite embryos were stained for Ac-tubulin and DAPI and subsequently mounted and imaged as described earlier. Zebrafish 14hpf embryos were stained with Ac-tubulin, Gamma-Tubulin and DAPI as per zebrafish whole mount immunofluorescence protocol. KV was dissected out and mounted on a slide. Mounted KVs were imaged on an Olympus Fluoview 1000 confocal microscope and Leica TCS SP8 AOBS inverted confocal microscope at x100 zoom and a z-stack images were taken for the entire vesicle. Mouse and fish images were processed using ImageJ with Simple Neurite Tracer utilised to measure each cilium from the base to the tip in 3 dimensions. After imaging, embryos were retrieved from the slides, washed with PBS and used for genotyping.

### Zebrafish KV Motile Cilia Live Imaging

Embryos from *cph* heterozygous fish incross were collected and placed in egg water at 28.5°C to develop until ∼14hpf. A mould designed to hold embryos was made using 2% agarose in egg water and left to set. Once embryos had developed to the correct stage, embryos with well-developed KVs were dechorionated and placed into the mould. Embryos were covered with egg water and placed on a Zeiss Imager M2 live imaging microscope. All embryos were imaged on a 100x water dipping lens at ∼150-200fps using Metamorph software. Embryos were then placed into separate wells of a 24 well plate coated with 2% agarose in egg water to stop the embryos sticking to the plastic. Recordings of each embryo were saved by well number. Embryos were left to develop until the mutant phenotype became apparent and recordings were correlated with the phenotype. Recordings were analysed using ImageJ and slowed to 15fps.

### RNA sequencing analysis

Total RNA was isolated from wild type and *Prpf8^N1531S^* mutant embryos at E10.5 using Trizol according to manufacturer’s instructions (Ambion). RNA was prepared from 3 embryos of each genotype. RNA quality was evaluated using TapeStation.

Paired-end RNA sequencing of E10.5 mouse embryo samples was performed on the Illumina HiSeq 3000 platform in the University of Leeds Next Generation Sequencing Facility. The FASTQ files were deposited into the Sequence Read Archive (SRA) under the accession number PRJNA690736. For RNA-Seq data, the adapter sequences were removed using Cutadapt (1.9.1) (87), and the filtering and trimming of sequences were done using PRINSEQ (0.20.4) (88). The clean reads were aligned to the UCSC mouse genome (mm10) (89) using STAR (2.5.1b) (90). The processing of BAM files was done using SAMtools (1.3.1) (91) and only uniquely mapped alignments were kept for further analysis. Read counts for protein-coding genes were generated through the featureCounts function provided by Rsubread package (1.22.0) (92) and RPKM values were calculated using rpkm function of edgeR package (3.14.0) (93). For those genes with RPKM >= 1 in at least one sample, a value of 1 was added to all RPKM values before the log2 transformation. Principal component analysis was carried out on the quantile normalised values and the first two principal components were plotted. We observed three expression outliers based on poor clustering: one from the wild type, one from the mutant correct looped, and one from the mutant reversed loop strain, which were excluded from downstream splicing analysis. Differential expression between groups of samples were conducted using DESeq2 package (1.12.0) (94) based on the raw counts for genes. Differential alternative splicing events were identified using rMATS (4.0.2) (55).

### Differential splicing analysis

Splicing events in RNA-seq data from the three wild-type day 10.5 embryos were compared against those in the three *Prpf8^N1531S^* mutant embryos using rMATS v4.0.2. Results were filtered to retain only splicing events exhibiting a difference in exon level inclusion (IncLevelDifference ≥ 0.2 or ≤ −0.2), and an FDR < 0.05.

### Gene ontology (GO) analysis

GO enrichment was performed using the PANTHER GO overrepresentation test (http://geneontology.org/). Genes harbouring exon skipping events with a difference in inclusion of ≥ 20% between the wild-type and *Prpf8^N1531S^* mutant embryos were evaluated against all *Mus musculus* genes in the GO Ontology Database. GO terms exhibiting a positive enrichment were retained. Genes present in the rMATS output were annotated with their respective GO annotations, retaining those annotations labelled as “biological_process” and with relationships to their genes of “involved_in” and “enables”.

### *Arl13b* isoform RT-PCR quantification

qPCR could not be utilised for isoform quantification as no primers or probe could be generated specifically for the “short” isoform of *Arl13b*. To quantify relative isoform levels, *Arl13b* RT-PCR was performed with a species-specific single primer set (Supplemental Table 4) that amplifies both the short and long isoforms. Gel electrophoresis was carried out to separate isoforms, with gels imaged on a Li-Cor Odessey and optical intensity quantified using Image Studio software with background gel intensity set to 0. The ratio of optical intensity of the “short” isoform to the “long” isoform was then calculated for each sample, and fold change calculated for the mutants as compared to wild type. Data are shown for at least three individual embryos of each genotype.

### Generation of Prp8^N1603S^ yeast and yeast splicing analysis

The N1603S mutation was introduced into pRS413-PRP8 by site directed mutagenesis (95). The pRS413-PRP8, pRS413-PRP8^N1603S^ or pRS413 plasmid was transformed into a haploid yeast strain where both the *CUP1* and *PRP8* genes were deleted and *PRP8* function was complemented with pRS416-PRP8. Growth on 5-fluoro-orotic acid (5-FOA) containing plates was used to select against and remove the complementing pRS416-PRP8 plasmid to leave the pRS413 plasmids as the sole source of *PRP8* and growth properties observed at 16°C, 30°C and 37°C. The resulting haploid strains with pRS413-PRP8 or pRS413-PRP8^N1603S^ were transformed with different *ACT1-CUP1* splicing reporters (53) and growth assayed on plates with increasing concentrations of CuSo_4_. Primer extension was carried out (53) to monitor the two steps of splicing from the *ACT1-CUP1* splicing reporters.

### ARL13B mammalian expression constructs

The long and short isoforms of human *ARL13B* were synthesised by GenScript and inserted as a N-terminal EGFP fusion protein into a mammalian expression plasmid with a CMV promoter. Sequence was confirmed by whole plasmid sequencing (Genewiz).

### hTERT-RPE1 cell transfections and imaging

#### Cell Culture

hTERT-RPE1 cells were cultured in DMEM/F12 media containing 10% (v/v) foetal bovine serum (FBS) and 2mM L-glutamine. Cells were split 1 in 10 using trypsin-EDTA when reaching 80-90% confluence. Cells were incubated at 37℃, 5% CO2.

#### Transfection

hTERT-RPE1 cells were transfected with *GFP*-tagged *ARL13B* long and short isoform plasmid using Lipofectamine^TM^ 3000 Transfection Reagent (Invitrogen), following the manufacturer’s instructions. The transfection medium was replaced with 0.2% FBS starvation medium after 24h and cells were incubated a further 24h-48h to induce cilia before use.

#### Immunofluorescence

hTERT-RPE1 cells on coverslips were fixed in 4% paraformaldehyde for 10 minutes at RT. Cells were then permeabilised with 0.3% Triton X-100 in PBS for 30 minutes at RT and blocked in SuperBlock blocking buffer (Thermo Fisher Scientific) for 1h at RT. Cells were incubated in the primary antibody (mouse anti-acetylated tubulin, Supplemental Table 5) diluted in ‘Solution 1 for primary antibody’ from SignalBoost Immunoreaction Enhancer Kit (Merck) overnight at 4°C before washing three times in 0.1% PBS-Tween20 and incubation in the secondary antibody (Alexa Fluor donkey anti-mouse 594 IgG (H+L) AffiniPure, Jackson ImmunoResearch) diluted in ‘Solution 2 for secondary antibody’ from SignalBoost Immunoreaction Enhancer Kit for 1h at RT. Cells were then washed three times in 0.1% PBS-Tween20 followed by incubation in DAPI for 5min. Coverslips were air dried before mounting with Prolong Gold antifade reagent (Thermo Fisher Scientific). Cells were imaged on a Leica TCS SP8 AOBS inverted confocal microscope. Images were processed with Fiji.

#### Zebrafish *arl13b* mRNA injections

Zebrafish *arl13b* long isoform was amplified by RT-PCR from zebrafish cDNA and cloned into PCS2+ xlt Vector using Hifi DNA assembly kit (New England Biolabs). Inverse PCR was used to generate the *arl13b* short isoform lacking exon 9. Plasmid inserts were sequenced (Genewiz). RNA was transcribed from a linearised plasmid using the mMessage Machine kit (Invitrogen). RNA was precipitated and stored at −80°C. RNA was diluted in DEPC-H_2_O to the desired concentration, and 0.5nl of 80ng/ml *arl13b* long or short isoform mRNAs was microinjected into zebrafish embryos at the single cell stage. Injected embryos were collected and placed in fresh egg water and left to develop at 28.5°C to the desired developmental stages. Embryos with green florescence were selected at 12hpf (using a LEICA M165 FC stereomicroscope) for use in further analysis.

#### *prpf8* splice blocking morpholino knockdown and rescue

Splicing blocking morpholinos targeting exon 7 of zebrafish *prpf8* (57) were synthesized by GeneTools (Supplemental Table 4). A 300nmol *prpf8* sbMO stock solution was made in sterile water to a final concentration of 1mM, and zebrafish embryos were injected with 0.5 nl of a 1:2 dilution of sbMO stock at the single-cell stage. For the rescue of *prpf8* morphants, after the injection of morpholinos, 0.5nl of *arl13b*-*GFP* long isoform or *arl13b-GFP* short isoform mRNA (200ng/ul) was injected into each morpholino injected embryo. Wild type embryos were injected with 0.5nl of a 1:2 dilution of Gene Tools standard MO as a control (Supplemental Table 4). Embryos were cultured in fish embryo medium (0.3g/L NaCl) until 12hpf. Embryos exhibiting green fluorescence by microscopy (LEICA M165 FC) were selected for further study. Embryos were incubated at 28.5°C until the desired developmental stages to observe the phenotype using brightfield microscopy (LEICA M80).

#### ARL13B Affinity purification-mass spectrometry Cloning and Vector Preparation

For GFP-trap affinity purification-mass spectrometry (AP-MS) analysis, the *ARL13B*_long and *ARL13B*_short isoforms were cloned into the CMV-EGFP-RGCC-SV40 poly(A)-NeoR/KanR vector (gifted by Prof. Ben Goult, University of Liverpool) by replacing the RGCC gene sequence with the respective *ARL13B* isoform sequences. Cloning was performed using SacI-HF and XmaI restriction enzymes (New England Biolabs, NEB, USA) and HiFi DNA Assembly, according to the manufacturer’s instructions. The control vector, expressing GFP alone, was generated by excising the RGCC sequence using MfeI-HF and BspEI restriction enzymes (NEB) and inserting a short filler sequence using HiFi DNA Assembly.

#### Transfection

1×10^6^ hTERT-RPE1 cells were seeded in 10cm plates one day before transfection. Transfection was performed using Lipofectamine 3000 transfection reagent (Invitrogen, L3000015) according to the manufacturer’s protocol. For each plate, 8µg of plasmid DNA was used. After 10 hours of transfection, the transfection medium was replaced with fresh complete medium to allow the cells to recover. At 24 hours post-transfection, the medium was changed to low-serum medium (0.2% FBS) for 24 hours to induce ciliogenesis before harvesting.

#### Cell Harvesting and Peptide Sample Preparation

hTERT-RPE1 cells expressing GFP-vector, GFP-ARL13B short isoform and GFP-ARL13B long isoform were harvested in RIPA buffer (Thermo Scientific, 89900, USA) supplemented with EDTA-free protease inhibitor cocktail (Roche, REF: 04693159001), PhosSTOP phosphatase inhibitor (Roche, REF: 04906837001), DNase (50 U/ml, Thermo Scientific, Cat: EN0521), and 2.5 mM MgCl2 (NEB, B0510A). Cell lysates were clarified by centrifugation. GFP pull-down, trypsin digestion, and peptide preparation for AP-MS analysis were performed according to the protocol provided with the ChromoTek iST Myc-Trap® Kit (Proteintech, Cat no: ytak-iST).

#### Mass Spectrometry Analysis

Samples were analyzed using a Thermo Exploris 480 mass spectrometer equipped with a FAIMS Pro interface and coupled to a U3000 nanoUPLC system. Each sample underwent a 1-hour LC-MS run, performed at the Biological Mass Spectrometry Facility, University of Manchester.

#### Data Analysis

Mass spectrometry data were processed using Fragpipe (v22.0), incorporating MSFragger (v4.1), IonQuant (v1.10.27), and DiaTracer (v1.1.5) within a label-free quantification (LFQ) and match-between-runs (MBR) workflow. MaxLFQ intensity values were exported from Fragpipe and analyzed using the R Shiny application “Manchester Proteome Profiler” (developed by Dr. Stuart Cain, unpublished). Proteins were identified as significantly enriched based on a fold-change threshold of >4 and an adjusted p-value cutoff of <0.05.

### Statistical analysis

Statistical analyses and the generation of graphs was performed on GraphPad Prism 10 for Mac, GraphPad Software, La Jolla, California, USA.

## Data Availability

RNAseq data can be accessed in the Sequence Reads Archive. Mass spectrometry data is available in the PRIDE database. Supplemental Movie Files are located on Zenodo with doi: 10.5281/zenodo.15482494.

## Acknowledgements

We thank Monica Justice for the *l11Jus27* mouse line and Graham Lieschke for the *cph* fish line. We thank Dominic Norris for advice on EM imaging of cilia, and for the *Nodal*, *Cerl2*, *Lefty1* and *Lefty2* mouse *in situ* hybridisation probe plasmids. We thank Megan Davey for advice on immunostaining of cilia. We thank Adam Hurlstone, Martin Lowe, Shane Herbert and Karel Dorey for advice on zebrafish experiments and access to equipment. Some computational data analysis work was undertaken on ARC4, part of the High-Performance Computing facility at the University of Leeds, UK. The authors wish to thank the staff in the University of Manchester FBMH Electron Microscopy Core Facility, Biological Mass Spectrometry Facility (RRID: SCR_020987), and Peter March in the BioImaging Core Facility for their assistance, and the Wellcome Trust for funding to support core facility equipment. We thank Stuart Cain for assistance with mass spectrometry data analysis and Xiaofan Xiong for uploading data to PRIDE. We thank the Biological Services Facility at the University of Manchester for mouse and fish husbandry. We thank the IMCB zebrafish facility for maintenance of zebrafish strains.

## Funding Statement

This work was supported by BHF grants PG/06/144/21898, PG/10/87/28624 and PG/18/28/33632 to KEH, Fight for Sight grant 5163/5164 to KEH, and funding from the Agency for Science, Technology and Research (A*STAR) of Singapore to S.R. KEH and BK acknowledge support from a British Heart Foundation Research Excellence Award (RE/24/130017). BK was supported by a BHF personal chair (CH/13/2/30154). DWM was supported by a University of Manchester-A*STAR Research Attachment Program (ARAP) Ph.D. fellowship. This work was supported in part by the Wellcome Trust [105610/Z/14/Z]. MB, LS and KM were supported by BBSRC DTP studentships awarded to the University of Manchester. MTG and DJS acknowledge EPSRC award EP/N021096/1.

## Author Contributions

FJ, MB, DM, WMSQ, CFR, GT, JZ, JL, KM, LAS, NA, DVL, EJRV, DW, MTG, BB, KS, AV, DS, KEH performed research. FJ, MB, DM, KEH wrote original manuscript draft. BK, MJH, JE, DS, CAJ, ROK, SR and KEH provided supervision and obtained funding. All authors read and edited the manuscript draft and approved submission.

## Competing Interests

No competing interests

## Materials and Correspondence

-KEH

## Notes

doi:10.5281/zenodo.15482494

### Competing Interest Statement

The authors have declared no competing interest.

### Summary of Updates

The title has been slightly altered from the first version to change "cilia function" to "cilia differentiation'. The funder Fight for Sight UK grant number 5163/5164 has been added to the funding statement. Typographical errors in Figure 6 legend have been corrected.

