## Supplemental Figures 1-6 for "The RNA splicing factor PRPF8 is required for left-right organiser cilia differentiation and determination of cardiac left-right asymmetry via regulation of *Arl13b* splicing"

**Supplemental Figures and Legends:**

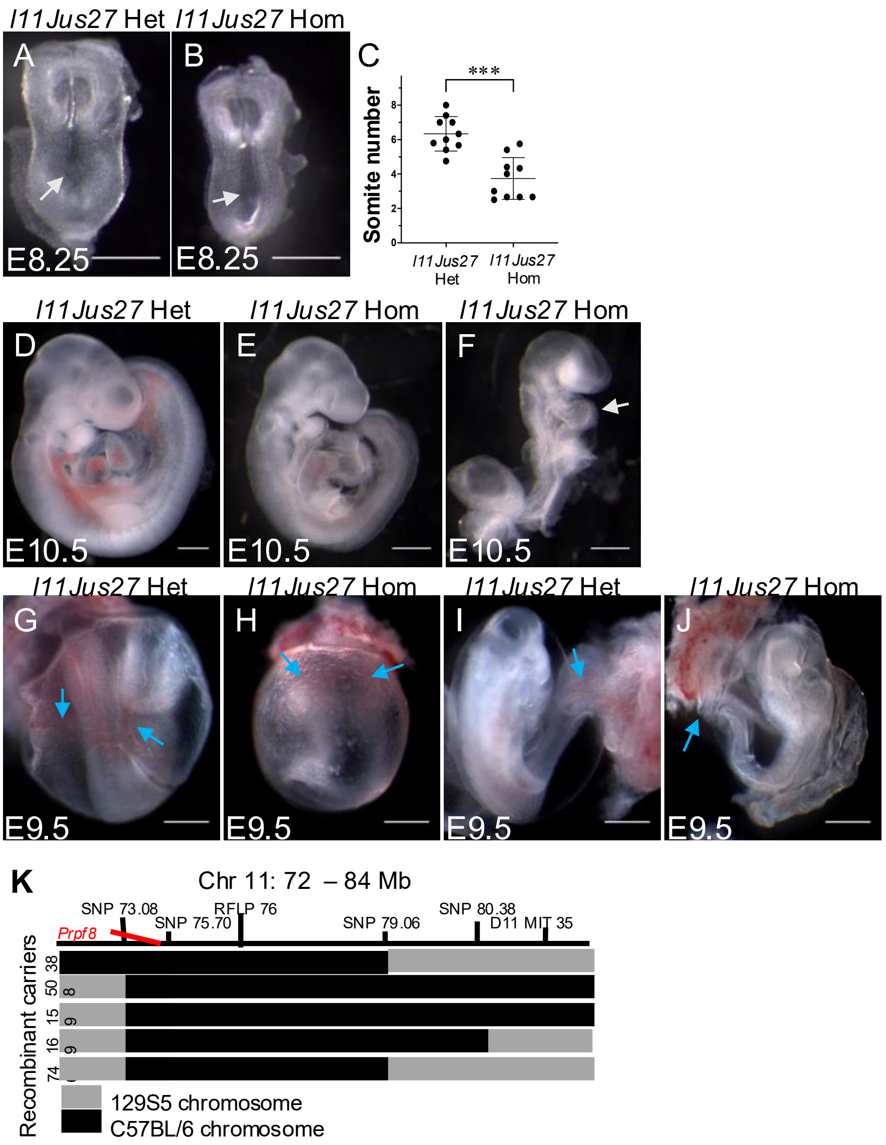


**Supplemental Fig 1. Morphological analysis of E8.5-E10.5 *l11Jus27* heterozygous and homozygous mutant embryos.** A) E8.25 *l11Jus27* heterozygous embryo. B) *l11Jus27* homozygous mutant embryo. Both embryos show normal headfold, somites, neural tube and node (arrow). C) Analysis of E8.5 embryos dissected between 12:00-13:00 reveals the average number of somites in *l11Jus27* homozygous embryos is significantly fewer than in littermate heterozygous controls ($p<0.0001$; two-tailed paired t-test; 10 pairs). Only dissections containing at least 2 *l11Jus27* homozygous embryos were analysed. D) E10.5 *l11Jus27* heterozygous embryo showing expected development. E) littermate *l11Jus27* homozygous embryo showing a mild phenotype with reversed cardiac looping and developmental delay. F) littermate *l11Jus27* homozygote showing severe delay in development. The embryo has not turned, chorioallantoic fusion has not occurred, the heart tube is underdeveloped and the gut and neural tube have not closed. G) *l11Jus27* heterozygous embryo showing prominent yolk sac blood vessels (blue arrows). H) *l11Jus27* homozygous embryo lacking mature yolk sac and umbilical vessels; only the primitive capillary plexus has formed (blue arrows). I) *l11Jus27* heterozygous embryo showing blood filled umbilical blood vessels (blue arrow). J) *l11Jus27* homozygous embryo lacking patent umbilical vessels (blue arrow). L= left, R = right. Scale bars = 0.5mm. K) Schematic diagram showing recombinant mapping of *l11Jus27* phenotype to a region of chromosome 11 containing *Prpf8*. The mutation is carried on the chromosome with the C57BL/6 genotype.

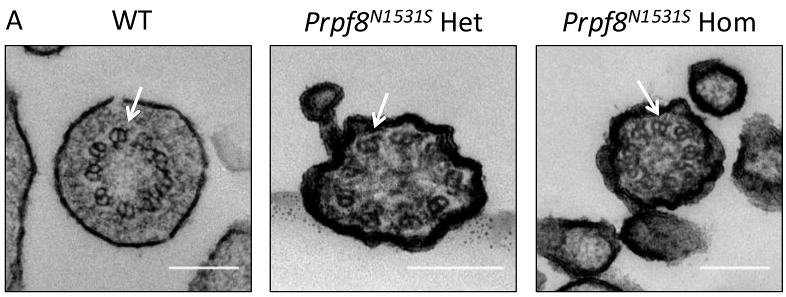


**Supplemental Fig 2. Transmission electron microscopy (TEM) of control and mutant node cilia.** A) TEM cross sections of nodal cilia were examined for defects in ciliary ultrastructure. *Prpf8*$^{+/+}$, *Prpf8*$^{N1531S/+}$ and *Prpf8*$^{N1531S/N1531S}$ nodal cilia all had dynein arms and the expected 9+0 configuration of microtubules. Shrinkage of the plasma membrane in the *Prpf8*$^{N1531S/+}$ and *Prpf8*$^{N1531S/N1531S}$ nodal cilia is likely a processing or fixing artefact (n=3 embryos for each genotype). Scale bar = 200nm.

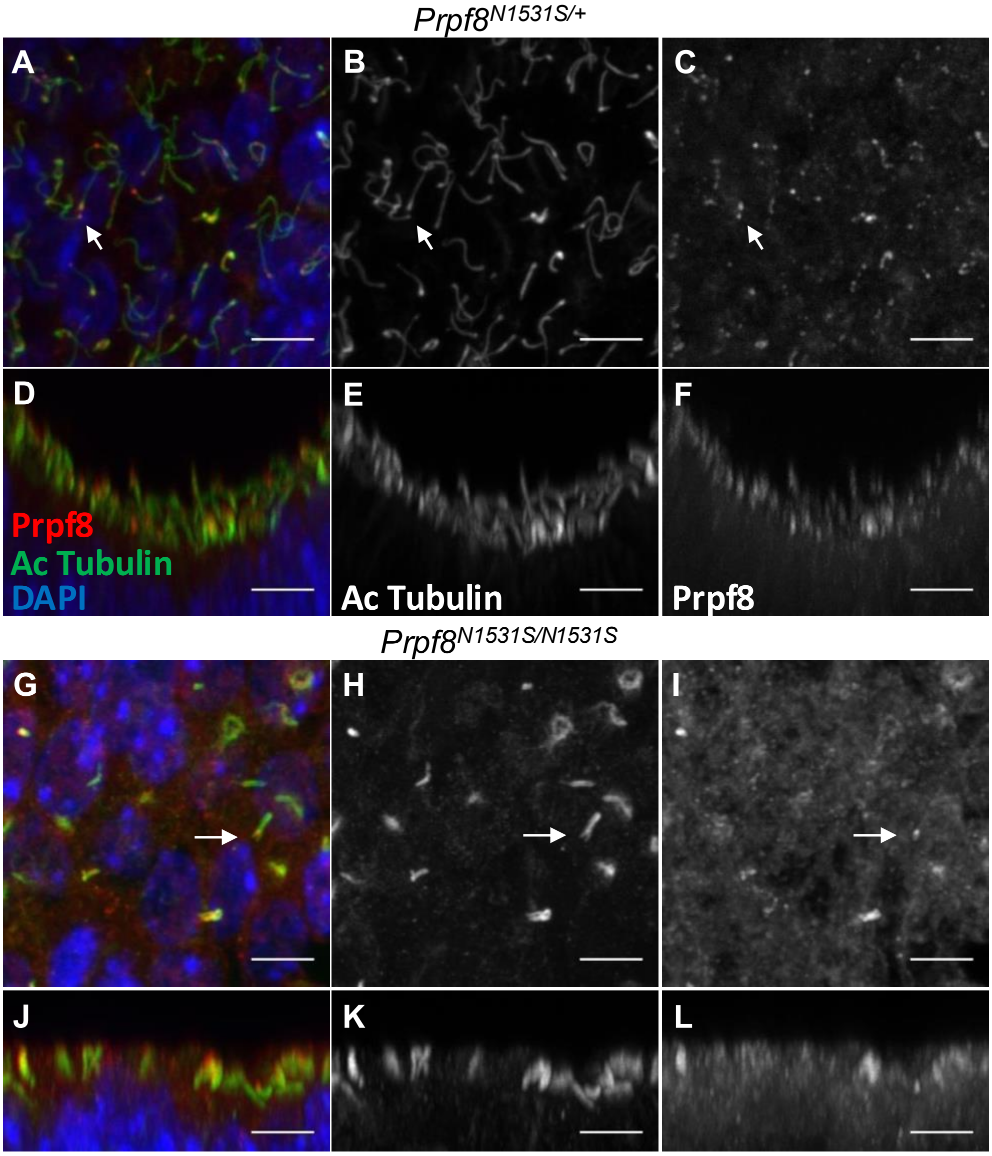


**Supplemental Fig 3. Localisation of PRPF8 in the node.** A-F) The staining pattern of anti-PRPF8 (red) in the embryonic mouse node was investigated via confocal microscopy; cilia were labelled with anti-acetylated tubulin (green) and nuclei were stained with DAPI (blue). There was no evidence of nuclear localisation of PRPF8 using these anti-PRPF8 antibodies. However, PRPF8 signal was detected at the presumptive basal body of the cilia. An example is indicated with an arrow. All images are maximum intensity projections of confocal z-stacks. A-C) Staining pattern of indicated antibodies in *Prpf8*$^{N1531S}$ heterozygous embryos (n= 5). D-F) Re-slice of A, B and C, respectively, showing side view of the node. G-I) Staining pattern of indicated antibodies on *Prpf8*$^{N1531S}$ mutant embryos (n= 7H). J-L) Re-slice of G, H and I, respectively, showing side view of the node, which is noticeably flatter compared to *Prpf8*$^{N1531S}$ heterozygous embryos. No difference in the staining pattern of either antibody can be discerned, though background is higher in the *Prpf8*$^{N1531S}$ mutant embryo. Scale bars = 5µm.

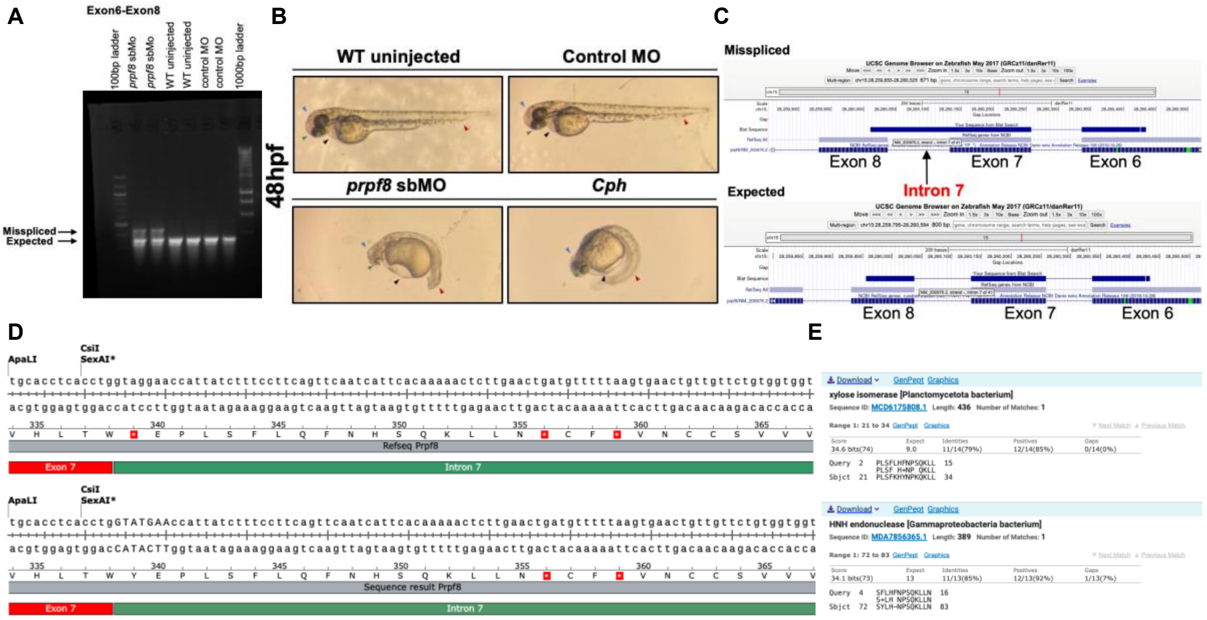


**Supplemental Fig 4. Validation of *prpf8* sbMO.** Splice-blocking morpholinos (sbMO) were designed to target exon 7 of *prpf8* because the exon length in bp is not a multiple of three, which is likely to create an out-of-frame protein translation sequence if incorrectly spliced, causing functional defects in prpf8. A) RT-PCR of sbMO injected fish using primers from *prpf8* exons 6-8 shows a larger product not detected in uninjected fish or fish injected with control MO. B) Morphology of WT uninjected fish and control MO injected fish was similar at 48hpf (top row). *prpf8* sbMO injected fish and cph mutant fish displayed similar morphological defects at 48hpf, including smaller heads (blue arrowhead), small eyes (green arrowhead), pericardial oedema (black arrowhead) and tail curvature defects (orange arrowhead; bottom row). C) Sequencing of the upper RT-PCR product present in sbMO injected fish confirmed that aberrant splicing occurs around exon 7, and that intronic sequence is included within the transcript. Note the *prpf8* reference transcript (blue) is shown from right to left on the chromosome. Similar results were seen with sbMOs targeting *prpf8* exon 24 (57). D) The predicted translation of the genomic reference sequence including intron 7 results in a stop codon at W338 (top). The sequence of the RT-PCR product from the sbMO fish differs from the reference fish genome sequence, however, the predicted translation of the aberrant intron-containing RT-PCR product results in the inclusion of 16 irrelevant amino acids prior to the inclusion of a termination codon and predicted truncation of Prpf8 at N355 (bottom). E) The 16 irrelevant amino acids generated in the sbMO fish aberrant transcript have some homology to bacterial enzyme sequences but are not homologous to Prpf8.

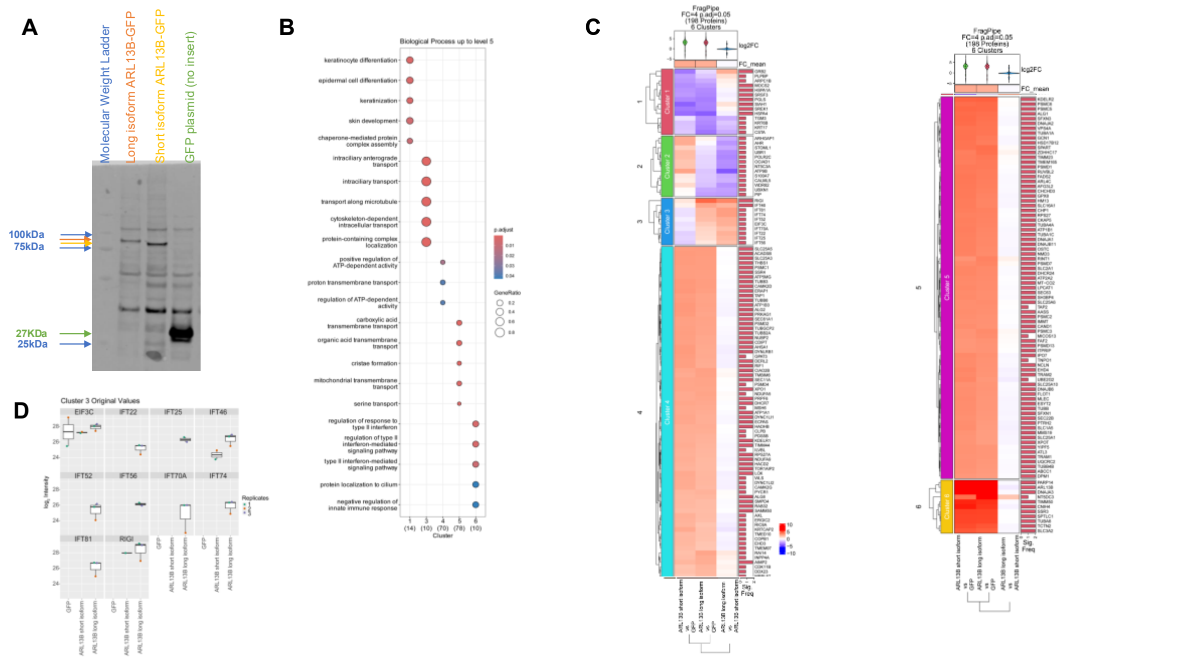


**Supplemental Fig 5. Mass spectrometry analysis of ARL13B long and short isoform proteins.** A) Western blot on RPE1 cell lysates detecting human ARL13B sequence with Nterminal GFP fusion. Molecular weight marker bands are identified in lane 1 with blue arrows. The long isoform (lane 2; orange arrow) produced a protein of higher molecular weight than the short isoform (lane 3; yellow arrow). The GFP plasmid without ARL13B sequence inserted produced a protein of the expected size for GPF (green arrow). B) Biological process annotations up to Gene Ontology (GO) Level 5 associated with proteins in Clusters 1, 3, 4, 5, and 6. Each biological process is represented by a spot, with the colour gradient indicating the adjusted p-value: red corresponds to lower (more significant) p.adjust values, and blue corresponds to higher (less significant) p.adjust values. The size of the spots represents the gene ratio, with larger spots indicating higher gene ratios. Both the colour and size legends are included in the figure to provide clear visual guidance for interpreting the enrichment significance and gene ratio values across clusters. C) Heatmap of proteins significantly enriched with a fold-change threshold greater than 4 and an adjusted p-value cutoff below 0.05. The visualization was generated using the "ComplexHeatmap" package and clustered into six groups using K-means clustering, grouping proteins with similar fold-change patterns. The x-axis indicates the comparisons: **ARL13B_long vs GFP**, **ARL13B_short vs GFP**, and **ARL13B_long vs ARL13B_short**. A hierarchical clustering tree is displayed on the left-hand side, further grouping proteins based on enrichment profile similarity. The colour gradient represents fold-change values, where **red** denotes upregulation (positive fold change) and **blue** indicates downregulation (negative fold change). The intensity of the colours corresponds to the magnitude of the fold change. The "Significant Frequency" column reflects the number of replicates in which each protein is significantly enriched. D) Box plots illustrating the distribution of non-imputed original MaxLFQ intensities for proteins enriched in Cluster 3. The x-axis represents the conditions, grouped by replicates and distinguished by colour coding, as indicated in the legend. The y-axis displays the log2-transformed intensity values.

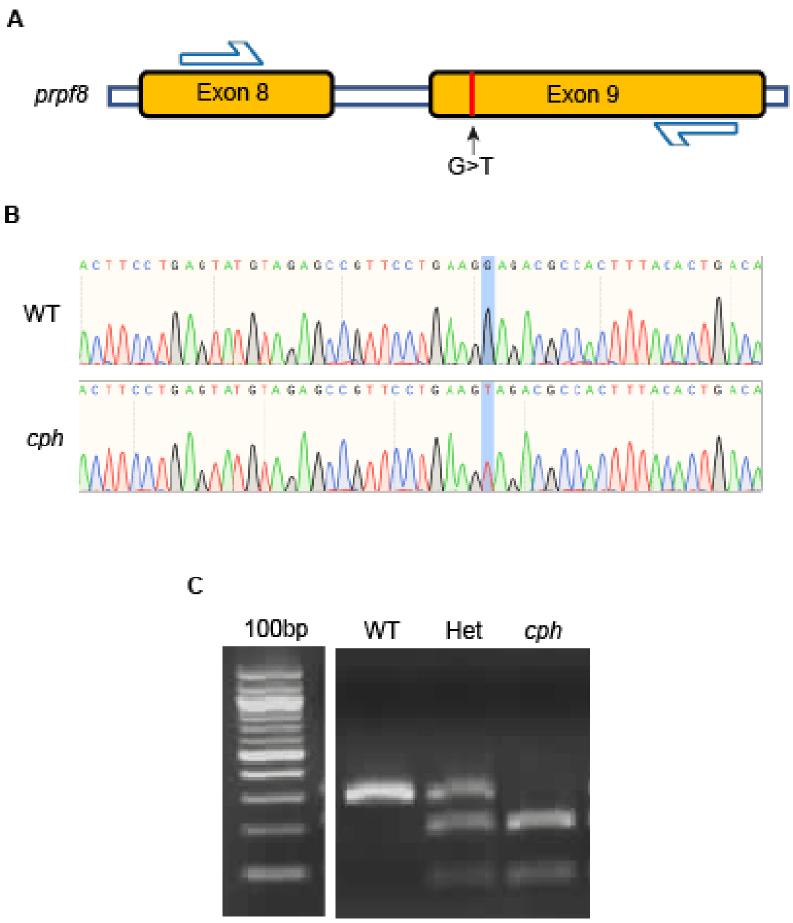


**Supplemental Fig 6. Genotyping of *cph* fish.** A) Schematic representation of mutated region of *prpf8* with location of genotyping primers added. B) Chromatograph from WT and *cph* embryos with mutated base highlighted. C) Gel electrophoresis of restriction digest using AccI enzyme on PCR reaction using primers from A.
